# Akt regulation of glycolysis mediates bioenergetic stability in epithelial cells

**DOI:** 10.1101/126771

**Authors:** Yin P. Hung, Carolyn Teragawa, Taryn Gilles, Michael Pargett, Marta Minguet, Kevin Distor, Briana Rocha-Gregg, Nont Kosaisawe, Gary Yellen, Joan S. Brugge, John G. Albeck

**Author notes:** These authors contributed equally (Yin P. Hung, Carolyn Teragawa).

## Abstract

Cells use multiple feedback controls to regulate metabolism in response to nutrient and signaling inputs. However, feedback creates the potential for unstable network responses. We examined how concentrations of key metabolites and signaling pathways interact to maintain homeostasis in proliferating human cells, using fluorescent reporters for AMPK activity, Akt activity, and cytosolic NADH/NAD^+^ redox. Across various conditions including metabolic (glycolytic or mitochondrial) inhibition or cell proliferation, we observed distinct patterns of AMPK activity, including stable adaptation and highly dynamic behaviors such as periodic oscillations and irregular fluctuations, indicating a failure to reach a steady state. Fluctuations in AMPK activity, Akt activity, and cytosolic NADH/NAD^+^redox state were temporally linked in individual cells adapting to metabolic perturbations. By monitoring single-cell dynamics in each of these contexts, we identified PI3K/Akt regulation of glycolysis as a multifaceted modulator of single-cell metabolic dynamics that is required to maintain metabolic stability in proliferating cells.

## Introduction

A central function of cellular metabolic regulation is to ensure an adequate supply of metabolites for bioenergetics and biosynthetic processes. To maintain metabolic homeostasis, cells utilize feedback loops at multiple levels in an integrated metabolic-signaling network. For instance, glycolysis is predominantly regulated by feedback control at the level of phosphofructokinase, which senses the availability of ATP and the respiratory intermediate citrate. Additionally, in response to ATP depletion, the energy-sensing kinase AMPK stimulates glucose uptake and suppresses energy-consuming processes (Hardie, 2008). The goal of these homeostatic pathways is to respond to bioenergetic stress by increasing or decreasing the appropriate metabolic fluxes to return the cell to a state with stable and sufficient levels of key metabolites. While bioenergetic stress can occur when any of a number of metabolites becomes critically limited, we focus in this study on the key metabolite ATP because of its broad importance as an energy source for cellular processes, and because AMPK activity can be used as a reliable indicator of low ATP:AMP ratios within the cell. We therefore use the term bioenergetic stress here to indicate a situation in which the concentration of available ATP is reduced, as indicated by AMPK activation.

Bioenergetic stress can result from a loss of ATP production, such as when nutrients become limited or metabolic pathways are inhibited by a pharmacological agent. Alternatively, ATP depletion can also result from an increase in ATP usage, such as when anabolic processes are engaged during cell growth. Because anabolic processes such as protein translation can use a large fraction (20-30%) of cellular ATP (Buttgereit and Brand, 1995; Rolfe and Brown, 1997), it is unsurprising that cellular proliferation and metabolic regulation are tightly linked (Gatenby and Gillies, 2004; Wang et al., 1976). Growth factor (GF) stimulation activates the PI3K/Akt pathway, which plays a key role in proliferation by stimulating both cell cycle progression and mTOR activity, leading to increased protein translation. Simultaneously, Akt activity promotes glucose metabolism by stimulating the activity of hexokinase (Roberts et al., 2013) and phosphofructokinase (Novellasdemunt et al., 2013) and translocation of glucose transporters (Glut1 and Glut4) to the cell surface (Sano et al., 2003; Wieman et al., 2007), while PI3K enhances the activity of hexokinase, phosphofructokinase, and aldolase to increase glycolytic flux (Hu et al., 2016; Inoki et al., 2012; Inoki et al., 2003).

The balance of anabolic and catabolic processes is particularly important in epithelial tissues, as they maintain the capacity to proliferate throughout adult life. Most cancers arise in epithelial cells (Koppenol et al., 2011) and involve a loss of both signaling and metabolic regulation (Gwinn et al., 2008; Vander Heiden et al., 2009). The AMPK and Akt pathways play key roles in this balance, intersecting through multiple crosstalk points and feedback loops to control both glucose metabolism (Fig. S1A) and protein translation at the level of mTOR. In principle, an optimal feedback response to an ATP-depleting perturbation would allow ATP levels to rapidly increase and stably restore ATP levels, while unstable responses such as continuing fluctuations or oscillations could be deleterious for the cell. However, a system with multiple feedbacks requires unavoidable tradeoffs in efficiency and robustness, and feedback increases the potential for instability (Chandra et al., 2011). Experimentally, such unstable metabolic responses have been observed in yeast (Dano et al., 1999; Ghosh and Chance, 1964) and in specialized post-mitotic mammalian cell types (Chou et al., 1992; O’Rourke et al., 1994; Tornheim and Lowenstein, 1973; Yang et al., 2008), confirming the potential for instability during metabolic adaptation. However, in epithelial cells, little is known about the kinetic relationships between signaling and metabolic activity that allow proliferation and other anabolic processes to proceed in step with energy production.

To understand the kinetics of homeostasis, single-cell data are needed because of the potential for metabolic state to vary even among genetically identical cells. Events that are asynchronous among cells, and subpopulations with differential behaviors, are not apparent in the population mean due to their tendency to “average out” (Purvis and Lahav, 2013). Until recently, dynamics in metabolism could only be measured effectively under conditions where fluctuations are synchronized across populations of cells, because biochemical techniques such as mass spectrometry provide broad measurement capabilities but reflect the population average rather than individual cells. However, advances in fluorescent reporters now enable real-time monitoring of metabolic and signaling dynamics in individual intact cells. Genetically-encoded fluorescent protein-based reporters have been designed to respond to specific metabolites by changes in their fluorescence output (Tantama et al., 2012; Tsien, 2005). As a result, metabolic and signal transduction states, including cytosolic NADH-NAD^+^ ratio (Hung et al., 2011; Zhao et al., 2015), glutathione redox potential (Gutscher et al., 2008), ATP-ADP ratio (Berg et al., 2009; Tantama et al., 2013), and AMPK activity (Tsou et al., 2011), can now be monitored in living cells.

In this study, we established a panel of proliferative epithelial cells expressing multiple fluorescent biosensors to enable detailed tracking of single-cell metabolic responses. We first used pharmacologic compounds to induce bioenergetic stress and to establish the range of cellular responses, finding that different forms of metabolic inhibition trigger strikingly different kinetics of adaptation and revealing conditions under which stable adaptation fails. We then used this framework to examine how metabolic adaptation functions in proliferating cells. We found that periods of bioenergetic stress occur throughout the normal cell cycle, and we identified a prominent role for PI3K/Akt regulation of glycolysis in mediating metabolic stability at the single-cell level.

## Results

### Fluorescent reporters enable single-cell imaging of metabolic and signaling dynamics

MCF10A mammary epithelial cells are dependent on GF stimulation for proliferation, can be stimulated to proliferate at different rates (Ram et al., 1995), and constitute a useful experimental system to examine the relationship between proliferation and metabolic kinetics. We generated MCF10A cell lines stably expressing a panel of genetically encoded fluorescent reporters for central components of the metabolic-signaling control network. First, the sensor AMPKAR2 (Fig. 1A) was constructed to monitor the activity of the energy-sensing kinase AMPK. Using the FRET-based AMPKAR biosensor (Tsou et al., 2011), we improved its dynamic range by using a brighter donor fluorescent protein mTurquoise2 (Goedhart et al., 2012) and an extended ‘EV’ linker (Komatsu et al., 2011). In live cells, AMPK activation was calculated based on a linear ratiometric method (Birtwistle et al., 2011), which we term ‘AMPK index’ (see Materials and Methods for details of activity calculations). We verified the sensing capability of AMPKAR2 by treating MCF10A-AMPKAR2 cells with the direct AMPK activator AICAR (Figs. 1B, 1C). Following AICAR application, all cells showed an immediate increase in the AMPK index and reached a steady state within 5 hours. When cultured in the absence of glucose, pyruvate, and glutamine for 24 hours, MCF10A-AMPKAR2 cells showed elevated AMPK index, which decreased abruptly upon glucose addition (Fig. 1D). Cells cultured in combinations of glucose, pyruvate, or glutamine displayed varying elevated levels of steady-state AMPK index (Figs.S1B, S1C), demonstrating that AMPKAR2 could monitor AMPK status across a range of physiological concentrations of nutrients in individual live cells.

**Figure 1.**
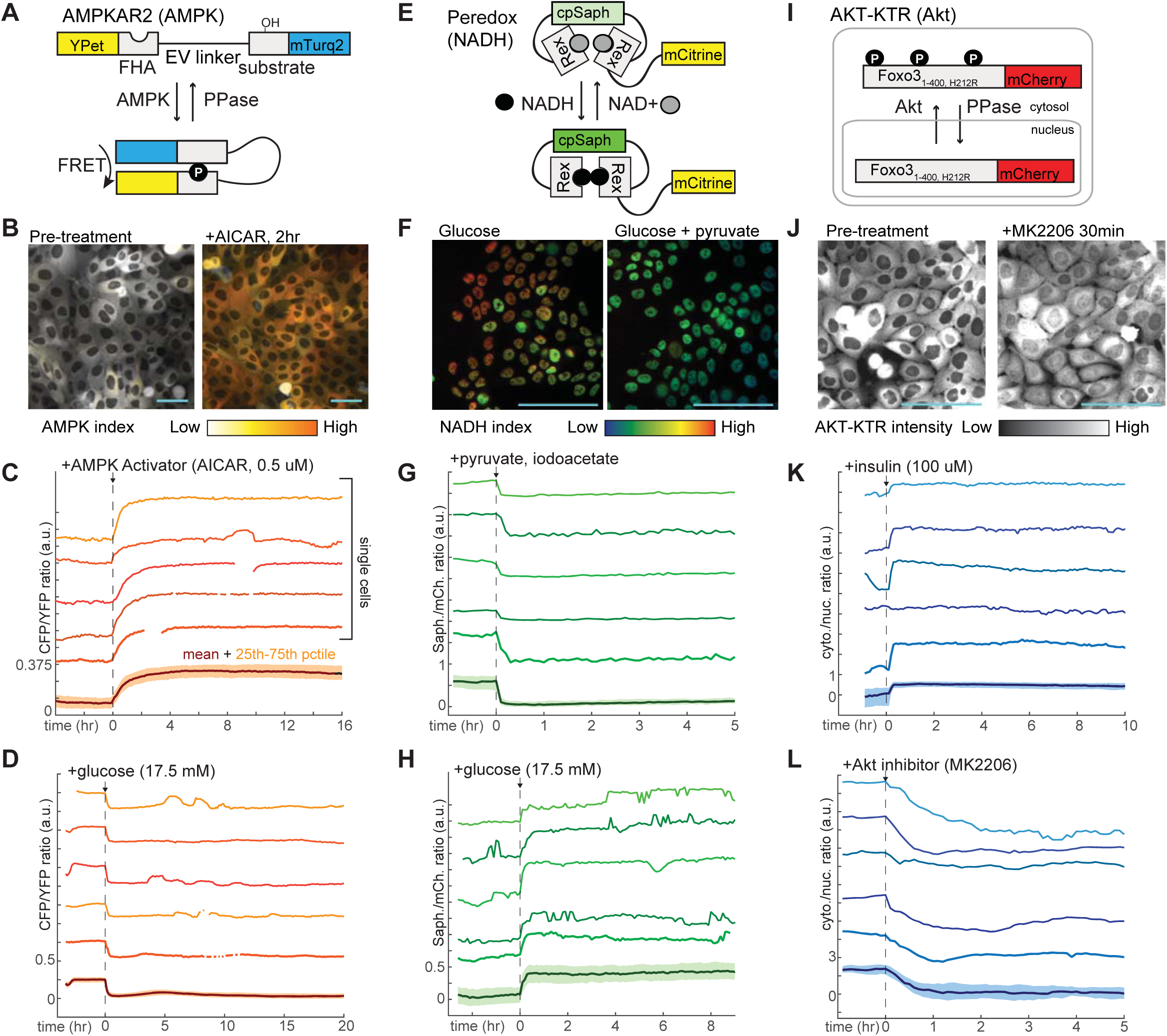
Design and validation of live-cell reporters for AMP-activated protein kinase activity (A-D), cytosolic NADH/NAD^+^ ratio (E-H), and Akt kinase activity (I-L). A,E, and I depict schematic diagrams for each of the reporters. B,F, and J show representative microscope images of MCF10A cells stably expressing each reporter. In images in B and F, reporter activity is represented as a pseudocolored image calculated as a ratio of the reporter components (see Methods for details of calculations). C, D,H, K, and L display individual and aggregate cell data for the indicated treatments. For each panel, the bottom-most profile represents the mean measurement for a population of >200 cells; the colored region around the mean indicates the 25th to 75th percentile range for the population. The five traces above the mean plot depict five representative individual cells, plotted at the same scaling as the mean. The media used in each experiment (prior to the indicated additions) were as follows: C – iGM (imaging-modified growth medium; see Methods); D – iGM lacking glucose; G – iGM2; H – iGM lacking glucose; K – iGM lacking insulin; L – iGM.

To assess the dynamics of the cytosolic NADH-NAD^+^ redox state, we utilized the fluorescent biosensor Peredox, which is based on a circularly-permuted green fluorescent protein T-Sapphire conjugated to the bacterial NADH-binding protein Rex (Hung et al., 2011). To maintain compatibility with red-wavelength reporters for dual imaging and to simplify cell tracking, we generated a nuclear-targeted Peredox fused to the YFP mCitrine (Fig. 1E). NADH is a major redox cofactor in glycolysis, which generates NADH from NAD^+^ via the glyceraldehyde-3-phosphate dehydrogenase (GAPDH) reaction in the cytosol. As NADH and NAD^+^ exchange freely between nuclear and cytosolic compartments, Peredox nuclear signal reports the cytosolic NADH-NAD^+^ redox state and serves as an indicator of glycolytic activity (Hung et al., 2011). Once normalized by the fused mCitrine signal to correct for variations in biosensor expression, Peredox nuclear signal is thus defined as the “NADH index.” To verify cytosolic NADH-NAD^+^ redox sensing, we exploited the lactate dehydrogenase reaction to interconvert between pyruvate and lactate with concomitant exchange between NADH and NAD^+^. MCF10A-Peredox cells treated with lactate or pyruvate in combination with iodoacetate as a glycolytic blockade exhibited rapid (<3 min) changes to reach maximal and minimal sensor responses, respectively (Figs. 1F, 1G, S1E), indicative of cytosolic NADH-NAD^+^ redox sensing (Hung et al., 2011). Consistent with previous data (Hung et al., 2011), steady-state NADH index increased with glucose concentrations (Fig. S1D), and Peredox could detect glycolytic dynamics in individual live cells (Fig. 1H). A control reporter with a mutation in the NADH binding site (Y98D) predicted to abrogate NADH binding failed to respond to the same conditions (Fig. S1E).

To track PI3K/Akt pathway activity, we constructed a reporter based on the Forkhead transcription factor FOXO3a. Akt phosphorylation of FOXO3a promotes its cytoplasmic retention; with low Akt activity, dephosphorylated FOXO3a translocates to the nucleus (Brunet et al., 1999; Tran et al., 2002). To monitor Akt activity, we thus fused a red fluorescent protein mCherry to a truncated FOXO3a gene in which transcriptional activity was abrogated to minimize any interference on endogenous gene transcription, a strategy previously shown to specifically report Akt activity (Gross and Rotwein, 2016; Maryu et al., 2016). We refer to this construct as AKT-KTR; the Akt activity indicated by its cytosolic-to-nuclear fluorescence ratio is referred to as the ‘Akt index’ (Fig. 1I). Upon insulin application following GF deprivation, MCF10A-AKT-KTR cells showed an abrupt increase in Akt index (Fig. 1K), consistent with the expected Akt stimulation. Conversely, pharmacologic treatment with the Akt inhibitor MK2206 induced an immediate decrease in Akt index (Figs. 1J, 1L), confirming that AKT-KTR could report Akt activity dynamics in individual live cells. We generated dual-reporter cell lines expressing both AKT-KTR and AMPKAR2, or AKT-KTR and Peredox, allowing Akt activity and metabolic status to be measured in the same cell. To track the relationship between proliferation and metabolic analysis, we also constructed a cell line expressing both AMPKAR2 and a reporter of S/G2 phase, GMNN-mCherry (Albeck et al., 2013; Sakaue-Sawano et al., 2008).

We used our reporter cell lines to establish the relationship between metabolic, signaling, and cell cycle parameters under different GF stimuli. Consistent with previous studies (Worster et al., 2012) insulin treatment induced the strongest activation of Akt index, while EGF produced a more moderate activation (Fig. 2A). Glucose uptake and NADH index were also highest in insulin-treated cells, intermediate in EGF-treated cells, and lowest in the absence of GFs (Figs. 2B, 2C), while the average AMPK index correlated inversely with Akt index (Fig. 2D). Because EGF stimulated proliferation more strongly than insulin (Fig. S2A), these metabolic parameters correlated poorly with proliferative rate. Together, these results suggest that the increased rates of glucose uptake and metabolism stimulated by Akt activity are the primary determinants of metabolic status under each GF. Accordingly, treatment of insulin-cultured cells with an Akt or PI3K inhibitor decreased glucose uptake (Fig. S2B) and NADH index (Fig. S2C) while increasing AMPK index (Fig. 2E). When NADH and AMPK index were tracked in individual cells, we found a high degree of variance over time within each cell, with some cells showing pronounced peaks and troughs (Figs. S2D, S2E). While this behavior appeared to correlate with certain GF conditions, we lacked a framework to quantify and interpret these dynamics; we therefore turned to defined metabolic perturbations as a tool to first establish basic homeostatic responses for single cells.

**Figure 2.**
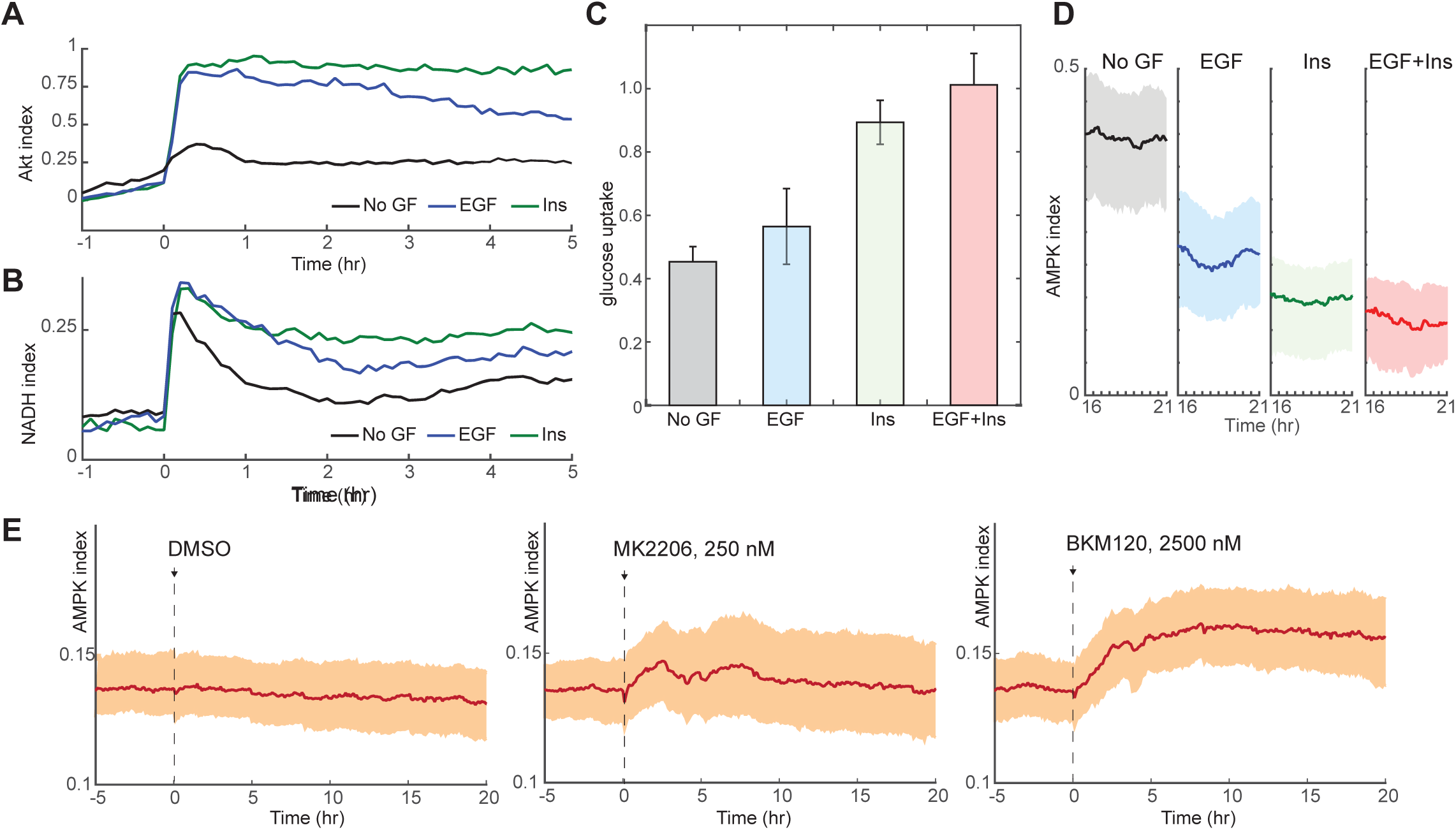
Glucose metabolism drives adaptation to the bioenergetic stress of proliferation. A. Mean Akt index, measured by Akt-KTR, following stimulation with EGF or insulin. Prior to imaging, cells were placed in medium lacking EGF or insulin. GFs were added at time 0. *N*=4, combined. B. Mean NADH index, measured by Peredox, following stimulation with EGF or insulin. Prior to imaging, cells were placed in medium lacking EGF or insulin. GFs were added at time 0. *N*=4, combined. C. Glucose uptake from culture medium by MCF10A cells. Glucose depletion from the medium was assayed immediately following a 2 hour period during which the cells were exposed to the indicated conditions. Bars are the average, and error bars the standard deviation, of 4 independent clones measured in triplicate in one experiment; results are representative of 3 total experiments run on different days. D. Mean and 25^th^-75^th^ percentile AMPK index for cells treated as in (A), recorded between 16 and 21 hours following treatment. *N*=5, representative. E. Mean AMPK index measurements for MCF10A cells cultured in medium containing glucose, glutamine, and insulin and treated at time 0 with either DMSO, 250 nM MK-2206, or 2.5 μM BKM120. *N*=4, representative.

### Fluctuating single-cell responses to metabolic challenges

To assess the range of individual cellular responses to specific bioenergetic challenges, we exposed MCF10A-AMPKAR2 cells to a panel of metabolic inhibitors, including oligomycin (an inhibitor of the mitochondrial F0/F1 ATPase), carbonyl cyanide m-chlorophenyl hydrazonesodium (CCCP, a mitochondrial proton gradient uncoupler), and iodoacetate (IA; an alkylating agent that inhibits the glycolytic enzyme GAPDH with minimal effects on other cellular thiols at <100 μM (Schmidt and Dringen, 2009)). As expected, each of these compounds rapidly raised the mean AMPK index in a dose-dependent manner in cells cultured in growth medium, confirming that both glycolysis and oxidative phosphorylation contribute to ATP production in proliferating MCF10A cells (Fig. 3, Supplemental Movies 1-3). However, each inhibitor induced strikingly different kinetics at the single-cell level. Notably, IA induced periodic oscillations of AMPK index, most evident at intermediate (5-10 μM) IA concentrations in which oscillations were sustained for as many as 50 cycles over 20 hours (Fig. 3A). These fluctuations of AMPK activity were not synchronized among individual cells and thus not apparent in the population average measurements. The asynchronous nature of these fluctuations argued against imaging artifacts or environmental fluctuations, which would affect all cells simultaneously. In contrast, oligomycin induced an immediate increase in AMPK index that peaked at ∼40 minutes but then fell, followed by a series of irregular pulses of AMPK activity ranging in duration from 1-3 hours (Fig. 3B).

**Figure 3.**
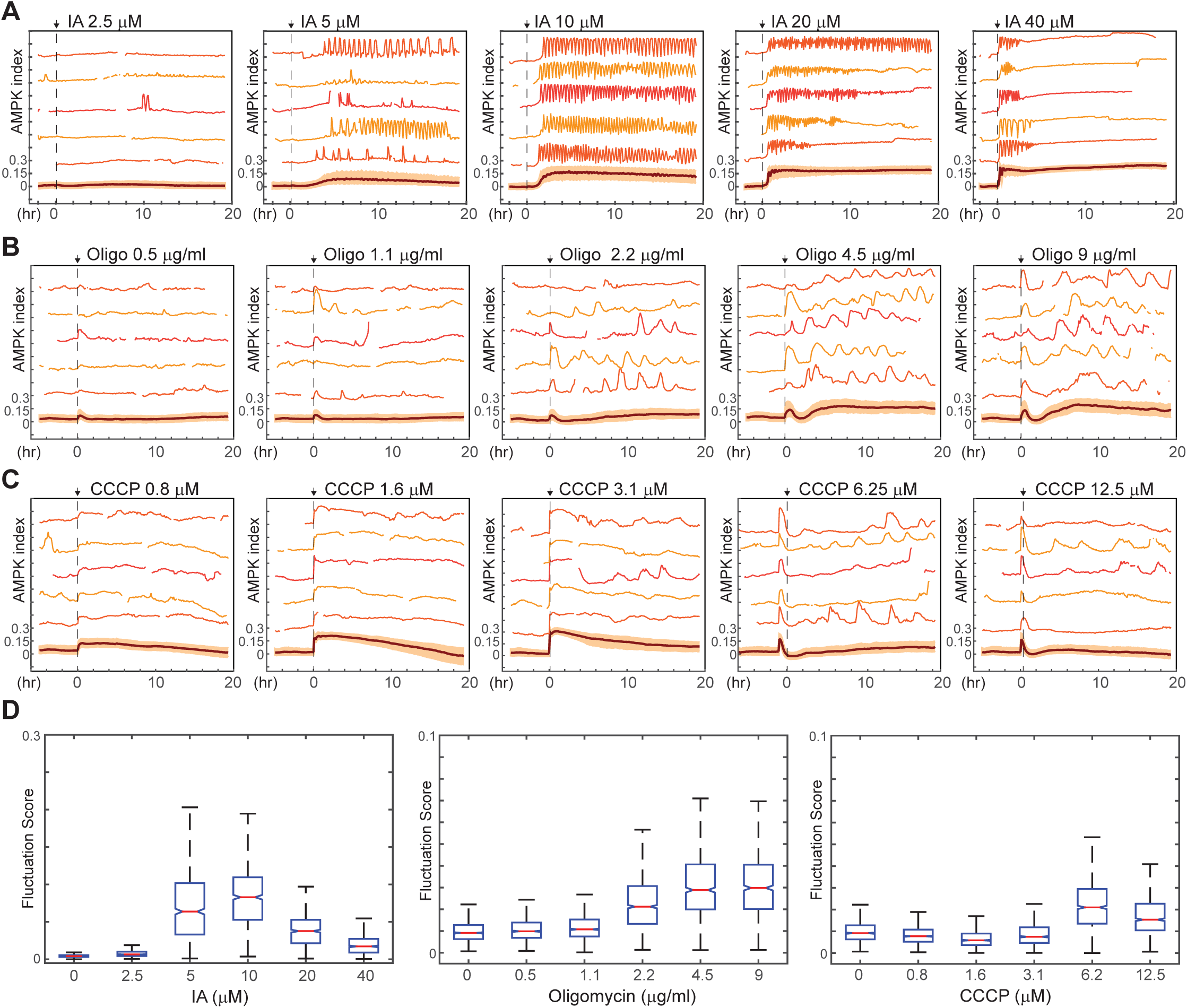
Kinetics of AMPK response to chemically induced metabolic stresses. A-C. Prior to imaging, MCF10A-AMPKAR2 cells were placed in iGM, and were then treated during imaging at time 0 with various concentrations of sodium iodoacetate (A), oligomycin A (B), or CCCP (C). Mean and representative single cell traces are shown as in
Fig. 1. *N*=3, representative. D. Fluctuation scores for each of the conditions shown in A-C, calculated as described in the Methods section. Scores were calculated for the period beginning 1 hour after treatment (to exclude the initial peak) and continuing through the end of the experiment.

Because pulsatile AMPK activities were a common feature of the single-cell response to multiple perturbations, we developed a “fluctuation score” to quantify the cumulative intensity of fluctuations for each cell over time (Fig. S3A and Methods section). Oligomycin and IA-treated cells showed significantly increased fluctuation scores relative to untreated cells (Fig. 3D). However, other perturbations resulted in different kinetic variations; for instance, at low and intermediate doses, CCCP induced a rapid increase in AMPK index, with a magnitude comparable to the other perturbations, that was maintained at a steady-state level for many hours with a low fluctuation score (Figs. 3C, 3D). Higher doses of CCCP, which are known to inhibit respiration, exhibited similar effects as oligomycin. We speculate that, at concentrations where CCCP acts only as an ionophore, a new stable steady state is reached due to ATP consumption by the F0/F1 ATPase working in reverse and pumping protons to maintain the mitochondrial electrochemical gradient; when proton flow is blocked by oligomycin (or potentially by high doses of CCCP), AMPK kinetics are determined by other processes, which we investigate below. Thus, bioenergetics and ATP levels exhibit distinct kinetics depending on the point of perturbation; each inhibitor induces a different reconfiguration of the metabolic network and pattern of feedback regulation.

Using AMPKAR2/GMNN reporter cells, we examined cell fates in response to each metabolic inhibitor (Supplemental Movie 4). IA-treated cells rapidly ceased cell cycle progression (Fig. S3B) and underwent lysis at varying times 12-24 hours following treatment. In oligomycin-treated cells, cell cycle progression slowed but still led to normal mitoses, and viability was unaffected. While 1 μM CCCP induced greater average AMPK index than oligomycin, neither cell cycle progression nor cell viability were altered. Thus, cellular responses to metabolic perturbations do not correlate with overall magnitude of bioenergetic stress. The observation that highly persistent dynamics are associated with more extreme changes in cell fate suggests that the kinetics of stress response may play an important role in determining cell fate.

### Temporally coordinated oscillatory dynamics in bioenergetics and signaling upon inhibition of glycolysis

To understand why cells fail to reach stable adaptation under some conditions, we focused first on the rapid oscillations triggered by IA treatment (Fig. 4A). The average period of these oscillations ranged from 18 minutes at 20-40 μM to 30 minutes at 5 μM (Fig. 4B). For IA at 10 μM or greater, the percentage of cells displaying oscillations (defined as 5 or more successive pulses with a spacing of 1 hr or less) was >95%; this percentage fell to <60% at 5 μM IA, and no oscillation was detected at concentrations of 2.5 μM or less (Fig. 4C). At intermediate concentrations (5-10 μM), IA-induced oscillations persisted for as many as 20 hours but typically ended with cell death (Supplemental Movie 4). Cells expressing Peredox or AKT-KTR treated with IA also exhibited oscillations in NADH index and Akt index, respectively, with kinetics similar to those seen in the AMPK index (Figs. 4D, 4E, Supplemental Movies 5 and 6). Oscillations were not observed when using the Peredox control reporter Y98D (Fig. S4A). To test whether the IA-induced oscillations in AMPK, Akt, and NADH indices were interrelated, we utilized AMPKAR3/AKT-KTR and Peredox/AKT-KTR dual reporter cells. AKT-KTR oscillations were tightly phase-locked with both AMPKAR2 and Peredox oscillations, that is, each cycle of AKT-KTR response corresponded to one cycle of AMPKAR2 signal and one cycle of Peredox signal (Figs. 4F, 4G). Phase locking was present in >90% of cells. In each cycle, peak signals of each reporter were phase-shifted relative to one another. In cells expressing Peredox and AKT-KTR, cycles initiated with a drop in NADH index that was followed approximately 0.25 cycles later by a drop in Akt index (Fig. 4F). In cells expressing AMPKAR2 and AKT-KTR, the initial decrease in Akt index coincided with an increase in AMPK index, and peaks of Akt and AMPK index remained shifted by 0.5 cycles thereafter (Fig. 4G). Based on these relative phase shifts, we constructed a composite diagram of the relationship between the three parameters (Fig. 4H). Thus, single-cell oscillations in PI3K/Akt activity, AMPK activity, and glycolytic NADH production were temporally coordinated, suggesting that feedback regulation tightly links these processes on the scale of minutes and leads to a persistent cycling of each pathway.

**Figure 4.**
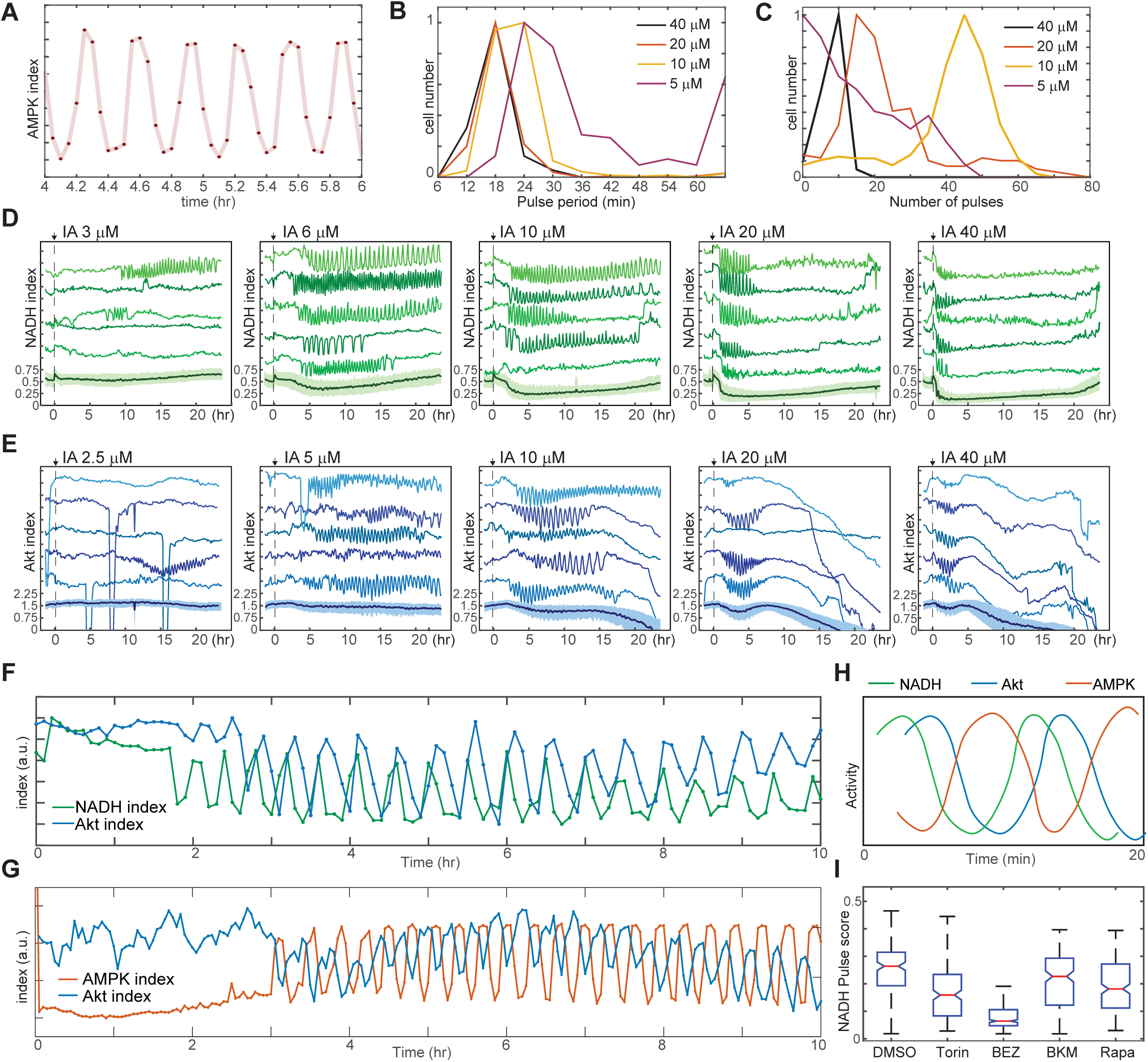
Linked oscillations in AMPK, Akt, and NADH indices triggered by inhibition of glycolysis. A. Expanded view of a representative region of oscillatory AMPK index for cells cultured in iGM and treated with 10 μM IA. B. Distribution of the average period of AMPK pulses for cells with 5 or more pulses, calculated as average peak-to-peak spacing following IA treatment as shown in Fig. 3A. Distributions represent measurements of >500 cells under each condition. C. Distribution of the number of pulses in AMPK index in each cell following IA treatment, measured as the largest series of detectable pulses spaced by 1 hour or less. D, E. Single cell measurements of NADH index (D) and Akt index (E) for the indicated concentrations of IA. Imaging medium was iGM lacking pyruvate, with IA added at time 0. *N*=4, combined. G. Simultaneous measurements of NADH and Akt indices (F) or AMPK and Akt indices (G) within the same cell. MCF10A cells expressing dual reporters were cultured and treated with 10 μM IA as in Fig. 4D and 3A, respectively. Cells shown were manually selected to best represent the phase relationship visible in the majority of cells. H. Diagram of approximate phase relationship between NADH, AMPK, and Akt indices derived from the data collected in (F) and (G). I. Pulse analysis of NADH index in IA-treated cells in response to inhibitors of PI3K/mTOR signaling. *N*=2-4, combined.

We hypothesized that IA-induced oscillations in AMPK activity and NADH/NAD^+^ ratio resulted from oscillations in glycolytic flux, triggered by feedback-driven increases in the entry of glucose into glycolysis upon GAPDH inhibition and flux reduction by IA. We therefore compared the fluctuation scores for IA-treated cells in the presence of varying extracellular concentrations of glucose and pyruvate. In the absence of glucose, IA treatment failed to induce oscillations in AMPK or NADH index (Figs. S4B, S4C). The incidence of oscillations, and the corresponding fluctuation score, increased with the extracellular glucose concentration in a dose-dependent manner, reaching a maximum at 4-5 mM, while the average period and amplitude remained essentially constant. In contrast, pyruvate alone, although capable of serving as an ATP source for MCF10A cells (Fig. S1B) was unable to sustain IA-induced oscillations (Fig. S4D). Pyruvate also had no effect on IA-induced AMPK index oscillations in the presence of glucose (Fig. S4D), although it rendered NADH index oscillations undetectable by lowering the resting NADH/NAD^+^ ratio (Fig. S4E). Together, these observations indicate that pyruvate does not fuel ATP production at a high enough rate to impact the rapid oscillatory changes during IA treatment. The data support the conclusion that these rapid oscillatory dynamics originate from changes in flux in glycolysis, with downstream metabolic processes playing little role.

Oscillations often arise in feedback systems in which there is a delay between induction of feedback and recovery of the feedback-controlled variables (Glass et al., 1988). The co-oscillation of Akt activity along with AMPK and NADH index suggests the involvement of a complex feedback structure involving PI3K/Akt, AMPK, and also mTOR (Fig. S1A) (Roberts et al., 2014; Yu et al., 2011). Consistent with this idea, multiple inhibitors of mTOR and PI3K activity suppressed IA-induced NADH oscillations (Fig. 4I). Suppression was most potent with BEZ-235, which inhibits PI3K, mTORC1, and mTORC2 activity, and strong but somewhat less potent with Torin1, which inhibits both mTORC1 and mTORC2. Both rapamycin, which inhibits mTORC1 alone, and BKM-120, which inhibits only PI3K, had more limited but still significant ability to block oscillations. Withdrawal of insulin from the growth medium, which simultaneously reduces Akt and mTOR activity and glucose uptake (Figs. 2C, S2B, and Worster et al., 2012), also resulted in attenuation of IA-induced oscillations (Fig. S4F). Taken together, the observation that oscillations are most potently suppressed when multiple candidate feedback controllers are inhibited supports the concept of a multi-tiered feedback control system. The phase relationship between the measureable variables in this system indicates the existence of delays between feedback activation and recovery of ATP and NADH (Fig. S4G) and supports a model in which the slowed flux through glycolysis due to IA treatment triggers a cyclic series of feedback events that drive regular oscillations (Fig. S4H).

### Modulation of glucose metabolism controls stable adaptation to mitochondrial ATPase inhibition

Oligomycin-induced fluctuations in AMPK differed from IA-induced oscillations in their longer time scale (2-6 hours), in their irregular nature, and in that they were not tightly associated with coordinated changes in NADH and Akt indices (Figs. S5A, S5B). To investigate the role of glycolysis in oligomycin-induced fluctuations, we examined the effect of glucose concentration in the absence of the alternate fuel sources glutamine and pyruvate (Fig. 5A, Supplemental Movie 7). At 0 mM glucose, the baseline AMPK index was high, and oligomycin treatment led to a small increase in AMPKAR index with no subsequent adaptation, followed by cell death in 100% of cells within 12 hours. In 17.5 mM glucose (the baseline concentration for MCF10A media), oligomycin induced a rapid initial pulse of AMPK activity and subsequent adaptation, with >75% of cells returning to baseline AMPK index within hours. Following this adaptation, cells displayed regular pulsatile dynamics in AMPK index, with an average period of ∼2.5 hours; the first two pulses of AMPK index were highly synchronous among cells, followed by gradual de-synchronization. As with the initial pulse, each burst of AMPK activity lasted 2-4 hours, suggesting that continuing oligomycin treatment induced ongoing bioenergetic challenges, which were nevertheless overcome by cells maintained at 17.5 mM glucose. At glucose concentrations of 3.4 mM and 1.7 mM, cells were unable to achieve full adaptation, with <25% and <10% of cells returning to baseline within 2 hours, and subsequent pulses in AMPK index were relatively dampened and prolonged. Thus, under high glucose levels, recovery of ATP levels occurs quickly and completely, but gives rise to recurring pulses of AMPK activity, suggesting that changes in the rate of ATP production by glucose metabolism via glycolysis generate these pulses.

**Figure 5.**
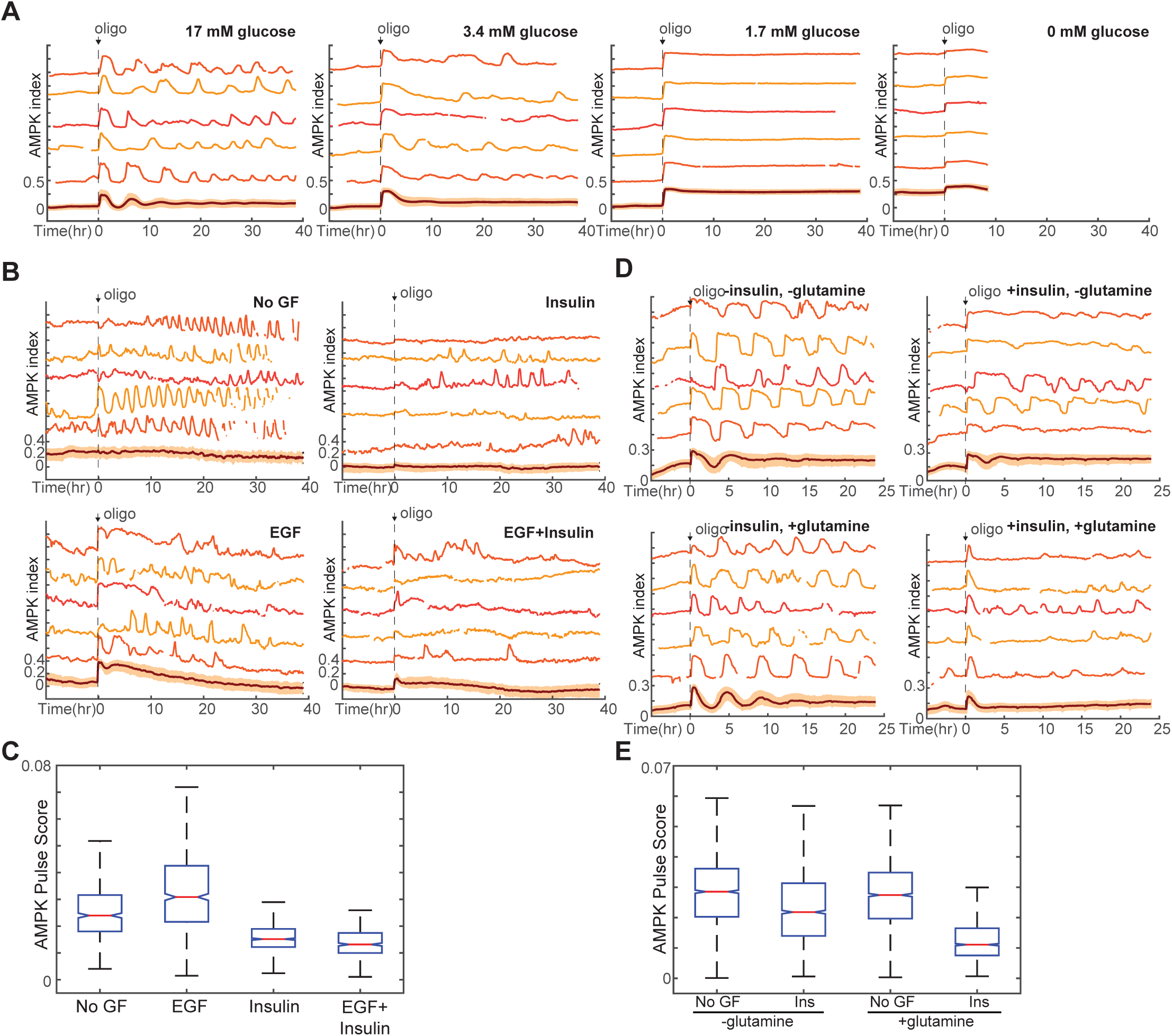
Requirement of glucose and glycolytic metabolism 828 for AMPK fluctuations. A. Single-cell measurements of AMPK index in response to oligomycin at different glucose concentrations. Prior to imaging, cells were placed in iGM lacking pyruvate and glutamine at different glucose concentrations, and were treated at time 0 with 1.8 μg/ml oligomycin. Measurements under 0 mM glucose are truncated due to cell death that began approximately 5 hours following oligomycin treatment. *N*=5, representative. B,C. Single-cell measurements of AMPK index in MCF10A cells in the presence of EGF, insulin, or both and exposed to 1.8 μg/ml oligomycin at time 0 (B). Quantification of fluctuation score for each condition is shown in (C). Prior to imaging, cells were placed in iGM lacking pyruvate and containing the indicated GFs. *N*=5, representative. D,E. Single-cell measurements of AMPK index in the presence of glutamine, insulin, or both (D); pulse quantification is shown in (E). Glucose was present at 17.5 mM in all conditions. *N*=2, representative.

We next examined how GF regulation of glucose metabolism impacts these dynamics (Fig. 5B). In the presence of glutamine, treatment with insulin strongly suppressed oligomycin-induced pulses in AMPKAR index, relative to non-GF treated cells. While EGF moderately increased the strength of pulses, co-treatment with both EGF and insulin led to suppression of pulses (Figs. 5B, 5C). This suppression was negated by co-treatment with Akt inhibitor, which strongly enhanced oligomycin-induced AMPKAR pulses (Fig. S5C). Similarly, in the presence of insulin, moderate inhibition of glycolysis with a dose of IA too low to independently stimulate oscillations resulted in amplification of oligomycin-induced AMPK pulses (Fig. S5D). Thus, the enhancement of glucose metabolism by insulin-mediated signaling is capable of attenuating recurrent ATP shortages following adaptation to oligomycin.

The pronounced and relatively regular nature of oligomycin-induced AMPK index oscillations in the presence of glucose alone (Fig. 5A), as compared to the more irregular pulses seen in complete growth medium (Fig. 3A), suggested that glutamine or pyruvate, which are present in complete medium, may also influence oligomycin-stimulated oscillations. While pyruvate had no effect on the kinetics of oligomycin response, the amplitude and regularity of pulsing were strongly attenuated when glutamine was present (Figs. S5E, S5F). When we performed the same experiment in the presence of different GF stimuli, we found that glutamine was required for the suppression of oscillations by insulin (Figs. 5D, 5E). However, regardless of the presence of insulin, cells cultured in the presence of glutamine without glucose failed to recover their AMPK index and died within 12 hours (Fig. S5F), suggesting that glutamine may play a role in supporting glucose metabolism, but cannot alone provide sufficient ATP in the absence of oxidative phosphorylation.

Altogether, these data suggest a model whereby, in the presence of high glucose, feedback regulation of glucose metabolism via glycolysis upon oligomycin treatment leads to recovery of ATP levels (Fig. S5G). However, this sharp increase in rate triggers negative feedback regulation of glycolysis, causing ATP levels to fall again (Fig. S5G, top). Insulin and glutamine counter these negative feedbacks, allowing glycolysis to continue at a high rate and thereby maintain high ATP levels (Fig. S5G, bottom). In contrast, in the presence of low glucose, oligomycin cannot stimulate a large enough increase in glycolysis to rapidly restore ATP levels, and negative feedback is not triggered, resulting in ATP levels that remain at lower, but stable, levels (Fig. S5G, middle).

### Akt-stimulated glucose uptake is required for bioenergetic stability in proliferation

Finally, we returned to the question of how GF stimulation and proliferation impact bioenergetic stability even in the absence of overt metabolic perturbations. We quantified the AMPK index fluctuation scores for GF-stimulated MCF-10A cells; we found that while AMPK index fluctuations under these conditions were less pronounced than in IA- or oligomycin-stressed cells, significantly more fluctuations occurred in non-GF- or EGF-treated cells relative to cells treated with insulin or a combination of insulin and EGF (Figs. 6A, 6B). To understand the basis for these fluctuations, we first used the Geminin cell cycle reporter to examine cell cycle-dependent differences in AMPK index. In EGF-stimulated cells in which AMPK fluctuations were most prominent, we compared the AMPK fluctuation scores between G0/G1 and S/G2/M phases of the cell cycle (Fig. 6C) and found that AMPK activity was significantly more pulsatile in G0/G1 cells. However, this moderate difference between cell cycle phases does not explain the overall effect of GFs on AMPK kinetics: compared to EGF-treated cells, insulin-treated cells are more likely to be in G0/G1 but have a lower probability of AMPK index fluctuations. We therefore next investigated the involvement of Akt by simultaneously monitoring both Akt and AMPK in dual reporter MCF10A-AMPKAR2/AKT-KTR cells. Analysis of fluctuations in both reporters on a cell-by-cell basis revealed a high frequency of inverse events, with a pulse in AMPK index mirrored by a decrease in Akt index (Fig. 6D). Cross-correlation analysis indicated that such inverse events were highly overrepresented in the population relative to their expected occurrences at random (Fig. 6E), suggesting that AMPK fluctuations may result at least in part from rises and falls in Akt activity and the associated rate of glucose uptake.

**Figure 6.**
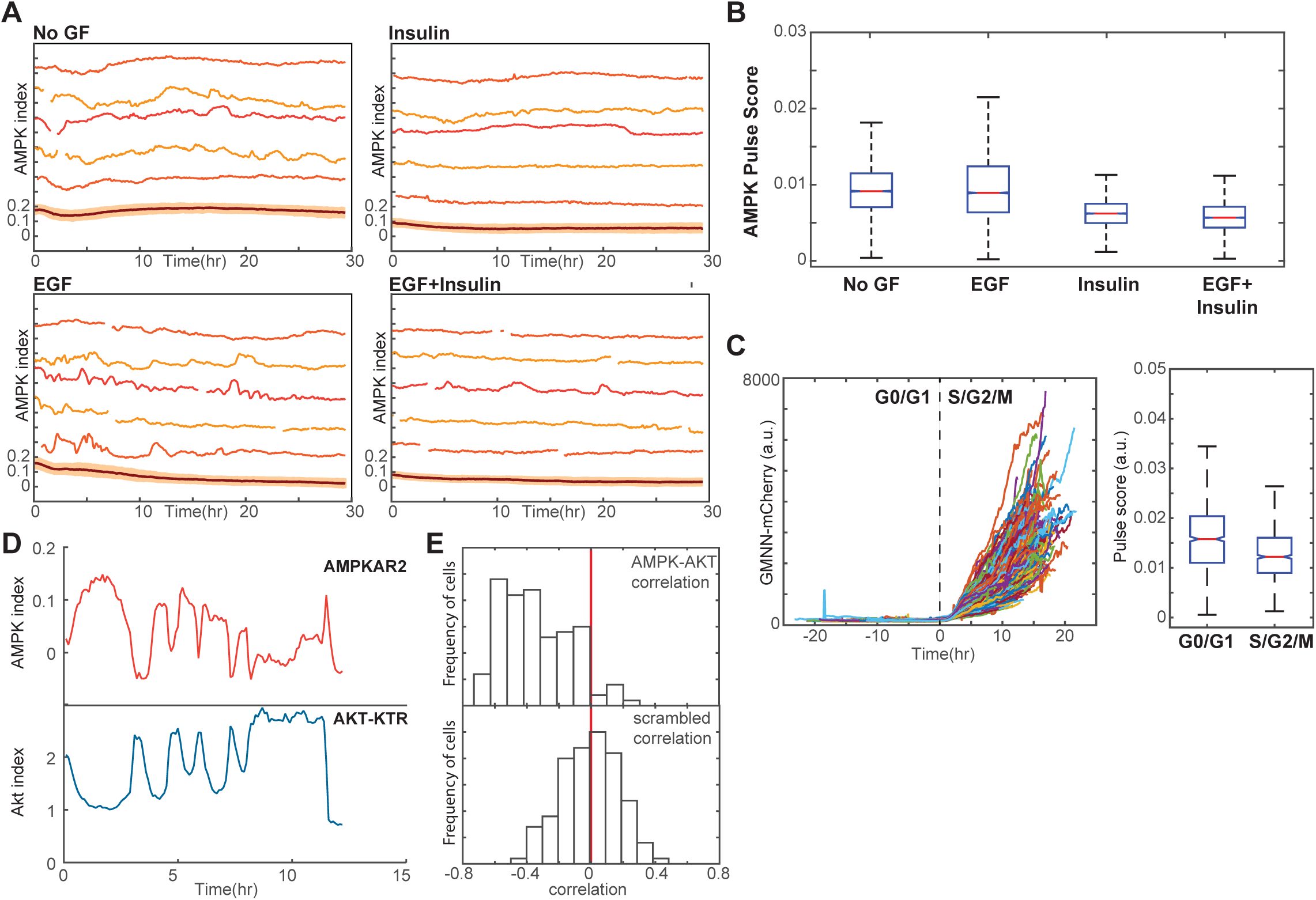
Linkage of AMPK and Akt activity fluctuations in the absence of chemical metabolic stresses. A,B. Single-cell measurements of AMPK index in MCF10A cells in the presence of EGF, insulin, or both. Prior to imaging, cells were placed in iGM lacking pyruvate and containing the indicated GFs. Quantification of fluctuation score for each condition is shown in (B). *N*=4, representative. C. Quantification of pulsatile behavior in G0/G1 relative to S/G2. Live-cell measurements of Geminin-mCherry and AMPK index were made in individual cells grown in the presence of EGF; cells were aligned by the time of G1-to-S transition (left). For each cell, we calculated the pulse scores during a 10-hour window preceding the induction of Geminin-mCherry to a 10-hour window immediately following induction (right). D. Single-cell measurements of AMPK index (AMPKAR2) and Akt index (Akt-KTR) in the same cell; culture medium was iGM lacking insulin and pyruvate. The cell shown was manually selected to display a prominent example of the anti-correlation trend visible in the population analysis in (E). E. Correlation analysis of AMPK and Akt indices. The time-dependent correlation between measured AMPK and Akt was quantified for each cells, and the distribution of individual correlation values is shown. A histogram centered around 0 indicates a population in which there is no systematic trend in the correlation, while skew to the left or right indicates a negative or positive trend in correlation, respectively. The bottom panel shows distributions of correlation values between Akt and AMPK for randomly chosen cells as a negative control.

We further tested the role of the PI3K/Akt pathway in bioenergetic stress using pharmacological inhibitors of this pathway. In cells growing in the presence of either insulin alone or a combination of insulin and EGF, treatment with either PI3K inhibitor (BKM120) or Akt inhibitor (MK-2206) increased the AMPK index fluctuation score (Figs 7A, 7B). Similarly NADH index fluctuations were increased with Akt, PI3K, mTOR, or dual PI3K/mTOR inhibitors (Fig. 7C). We conclude that PI3K/Akt signaling plays an active role in suppressing fluctuations in bioenergetic stress in cells at both high and low proliferation rates.

**Figure 7.**
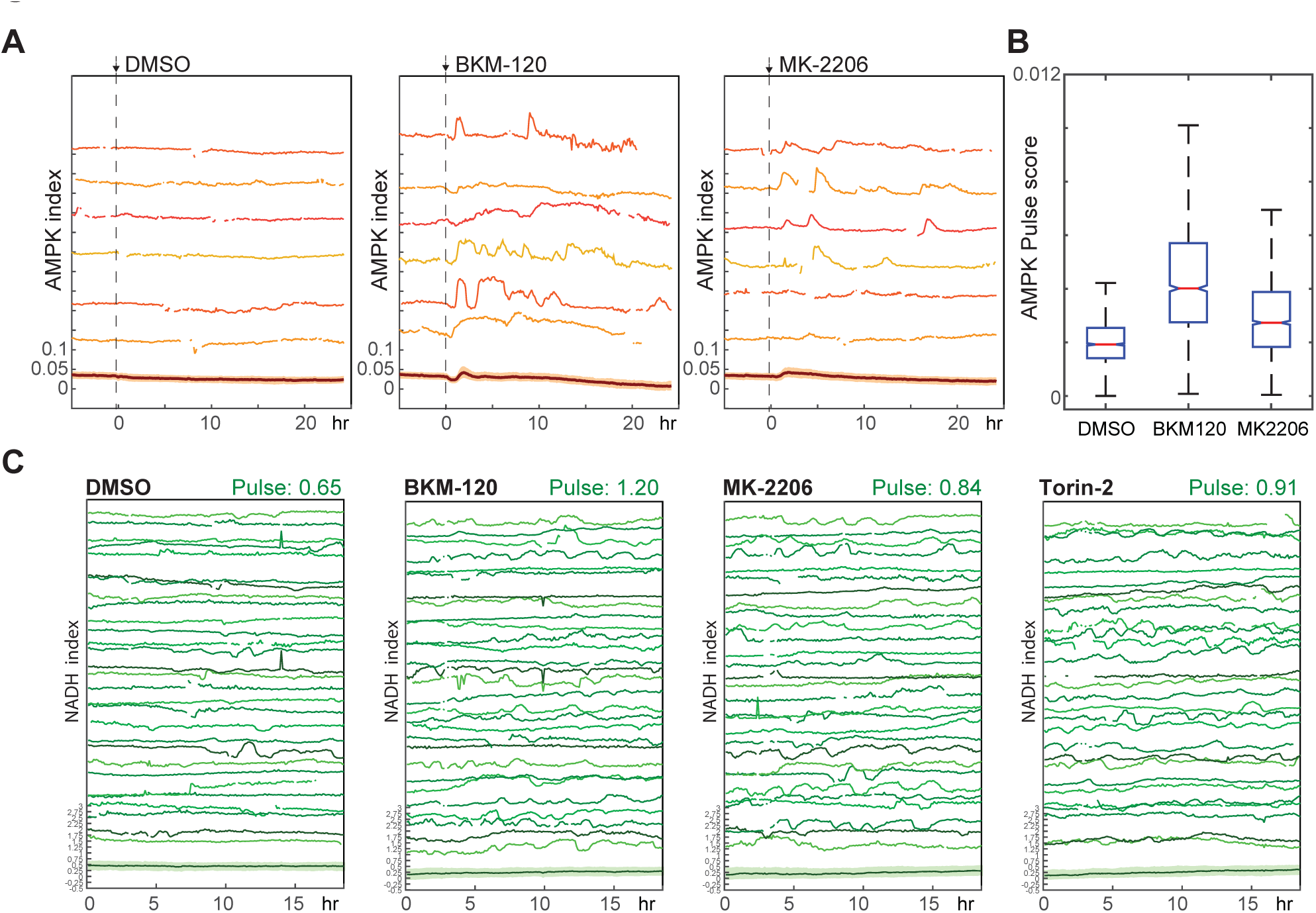
Suppression of spontaneous bioenergetic fluctuations 860 by Akt/PI3K signaling. A,B. Pulse analysis of AMPK index in MCF10A cells imaged in iGM and treated with BKM-120 (2.5 μM) or MK-2206 (250 nM) at time 0. D. Pulse quantification for the conditions in (B). *N*=4, representative. C. NADH index in MCF10A cells expressing Peredox and cultured in iGM2 with the indicated inhibitors (added 5 hours prior to the beginning of the experiment). Due to high variability from cell to cell, a larger number (30) of randomly chosen traces are shown for each experiment to better display the range of behaviors. Median pulse scores are shown in green above each plot*. N*=5, combined.

## Discussion

### Balancing metabolic adaptability and stability at the single cell level

The kinetics of metabolic homeostasis at the organismal level are known in detail (for example, blood glucose clearance rates are well characterized and widely used as diagnostics), but metabolic homeostasis at the single cell level has remained largely unexplored. Given the extensive interconnections between metabolic regulation and GF signaling at the level of Akt, mTOR, and AMPK, intricate metabolic dynamics in single cell physiology have long been postulated (Plas and Thompson, 2005) but never explored in detail. Here, using a panel of fluorescent biosensors for key metabolic regulators, we show that individual cells experience frequent deviations in bioenergetic and signaling parameters, both during proliferation and in response to metabolic challenges. As fluctuations in ATP and NADH availability can influence functions such as DNA synthesis and gene expression, understanding these metabolic dynamics, rather than simply average or baseline concentrations, will be crucial in developing an integrated model for the control of cellular metabolism and growth.

Feedbacks in metabolic control enable the cell to maintain adequate levels of key metabolites under non-ideal circumstances. Because ATP plays a central role in providing energy for many essential cellular processes, even short lapses in availability can potentially compromise cellular function and viability; it is likely that evolution has selected for feedback kinetics that rapidly reverse any decrease in ATP to prevent levels from falling dangerously low. Consistent with this idea, we find that cells provided with different fuel sources (glucose, glutamine, and pyruvate) are able to adapt and maintain steady levels of AMPK activity with few fluctuations, albeit at different set points that depend on the fuel source (Figs. S1B, S1C). However, optimization for such rapid and efficient adaptation comes with the potential that for certain conditions stable adaptation cannot be achieved, and unstable (e.g. oscillatory) responses result (Chandra et al., 2011). In terms of nonlinear dynamics, such responses occur in limited regions of parameter space near unstable fixed points. Accordingly, we find that epithelial cells can be forced into persistent oscillatory behavior within certain intermediate conditions. For example, IA-induced oscillations are most persistent at intermediate doses of 10-20 μM; at lower doses, cells successfully adapt after brief cycling, while, at higher doses, cells simply fail to adapt and remain at a high level of AMPK activity and low NADH. The conditions where we observe unstable behavior – including inhibition of lower glycolysis or mitochondrial ATP production, or culture in the complete absence of insulin-stimulated glucose uptake – likely represent situations that are unusual under normal physiological function; however, such conditions may occur under pathological circumstances, including mutations of metabolic enzyme genes or pharmacological or toxic compounds that impair metabolic function, and may be important for the understanding of tissue function in these cases. In the epithelial cells examined here, metabolic oscillations coincide with decreases in cellular function ranging from slowed proliferation to cell death, suggesting that the oscillations represent a deleterious side effect of feedback control. Nevertheless, it has also been proposed that sustained glycolytic oscillations could play an important role in fine-tuning certain cellular functions, such as insulin secretion in pancreatic beta cells (Goodner et al., 1977).

### Glycolysis and insulin signaling in the control of metabolic stability

Our results point to a central role for glycolysis in mediating metabolic stability. Despite its relative inefficiency in ATP yield per molecule of glucose, glycolysis can, at least under certain conditions, produce ATP at a faster rate than oxidative phosphorylation if sufficient glucose is available; this ability is best documented in muscle cells during anaerobic activity but could conceivably extend to other situations such as hypoxic cells within a tumor (Liberti and Locasale, 2016). Our results also implicate insulin and the PI3K/Akt pathways as controlling factors in bioenergetic stability, consistent with their stimulatory effect on glucose uptake and glycolytic flux. The data presented here suggest that the capacity for rapid ATP production by glycolysis can play both positive and negative roles in bioenergetic stability. For example, insulin enhances the occurrence of IA-induced oscillations (Fig. 4I), but has a suppressive role for oligomycin-induced oscillations (Fig. 5C). In our oscillation models (Figs. S4G, S4H, and S5G), this difference is consistent with the configuration of the network in each case. In the case of IA, where a bottleneck is imposed between upper and lower glycolysis, higher glucose input stimulated by insulin would increase both the maximum rate for ATP production and the strength of negative feedback, but flux from upper to lower glycolysis would remain limited by the inhibitor, leading to stronger oscillations. In the case of oligomycin, increasing the flux through glycolysis to its maximum rate enables the pathway to produce sufficient ATP in the absence of oxidative phosphorylation, preventing negative feedback from AMPK to initiate oscillations. Thus, while glycolysis is the preferred route to quickly restore ATP levels when they fall, the resulting rapid changes in ATP and other metabolites may also facilitate oscillatory behavior, as slower regulatory processes attempt to catch up to the increased glycolytic flux.

Previous studies of glycolytic oscillations have measured periods ranging from several seconds to twenty minutes (Chou et al., 1992; O’Rourke et al., 1994; Tornheim and Lowenstein, 1973; Yang et al., 2008). Oscillations have been recapitulated in isolated extracts of both yeast and mammalian myocytes, indicating that the core glycolytic enzymes alone are sufficient, with the allosteric regulation of PFK playing a central role. Our observations differ from these studies in the longer period of the oscillations (20-30 minutes in the case of glycolytic inhibition by IA; 3-6 hours in the case of mitochondrial inhibition by oligomycin), as well as in implicating a role for Akt and AMPK in oscillations. We speculate that in the epithelial cells examined here, a core glycolytic oscillator becomes entrained through feedback connections to these additional regulatory pathways that are central to growth and homeostasis in this cell type. Our data also suggest that metabolic oscillations may be a wider phenomenon than previously thought, as we demonstrate their occurrence in cells not typically considered highly metabolically active, and also find that they may occur with heterogeneous phasing that makes oscillations impossible to detect without single cell methods. The tools developed here will be of use in detecting and analyzing similar oscillations in other cell types and conditions.

### Implications of energetic stability in GF signaling, carcinogenesis, and pharmacotherapy

Given that over 90% of human tumors arise in epithelial tissue and that abnormal cell proliferation underlies carcinogenesis, understanding metabolic requirements for proliferating epithelial cells can have profound implications in oncology research. Our findings offer a potential explanation for the metabolic advantage conferred by aerobic glycolysis in tumors and proliferating cells. Existing hypotheses for why aerobic glycolysis is common in proliferating cells include rapid ATP generation by glycolysis (though less efficient than oxidative phosphorylation), as well as increased fluxes to glycolytic intermediates for biosynthesis (Sullivan et al., 2015). We find that in the presence of EGF, where ATP and NADH are low but proliferative rate is high, cells display increased bioenergetic instability and sensitivity to inhibition of oxidative respiration, which can be reversed by insulin-mediated activation of Akt. In the context of tumor microenvironments with fluctuating nutrient and oxygen supply, such instability is likely deleterious and may create selective pressure for genetic alterations to enhance glycolysis, such as activating mutations in the PI3K/Akt pathway, which are among the most frequent mutations across all cancer types. Investigating the effects of oncogenic mutations on metabolic stability may thus be important in developing therapies that target the altered metabolism of tumor cells.

## Methods

### Reporter construction

Peredox-mCitrine-NLS was constructed by replacing mCherry with mCitrine (Shaner et al., 2005) in pMSCV-Peredox-mCherry-NLS (Hung et al., 2011). The negative control with abrogated NADH binding was constructed by introducing the mutations Y98D in both subunits of Rex and I189F in the first subunit of Rex in pMSCV-Peredox-mCherry-NLS. AKT-KTR was constructed by fusing the N-terminal domain (amino acid residues 1–400) of human FOXO3a DNA binding mutant H212R (Tran et al., 2002; from Addgene) with a C-terminal mCherry in the retroviral pMSCV vector. AMPKAR2-EV was constructed by modifying the linker between the CFP and YFP in AMPKAR (Tsou et al., 2011) with an expanded EEVEE linker (Komatsu et al., 2011), and replacing the CFP and YFP fluorophores with mTurquoise2 and YPet, respectively; a PiggyBAC transposase-mediated delivery system (Yusa et al., 2011) was used to minimize recombination between CFP and YFP.

### Reagents

Reagents were from Sigma unless noted. Iodoacetate, lactate, pyruvate, and cycloheximide stocks were prepared in water. Rotenone, oligomycin A, and AICAR were dissolved in DMSO. BEZ235 (Axon Medchem), Torin-1 (Tocris Bioscience), Torin-2 (Selleck), BKM120 (Axon Medchem), GDC0941 (Axon Medchem), LY294002 (Sigma), Gefitinib (Axon Medchem), MK2206 (Selleck), PD 0325901 (Calbiochem and Selleck), and Rad001 (SU2C PI3K Dream Team Mouse Pharmacy, which obtains compounds from Shanghai Haoyuan Chemexpress; (Elkabets et al., 2013)) were dissolved in DMSO. For GF titration, epidermal growth factor (EGF; Peprotech) and insulin (Sigma) were diluted in PBS and added at indicated concentrations.

### Cell Culture and Media

Human mammary epithelial MCF10A cells and the clonal derivative 5E (Janes et al., 2010) were cultured as previously described (Debnath et al., 2003). The MCF10A full growth medium consisted of Dulbecco’s modified Eagle’s medium (DMEM)/F-12 (Life Technologies 11330), supplemented with 5% horse serum (Life Technologies), EGF (20 ng/ml), insulin (10 μg/ml), hydrocortisone (0.5 μg/ml), cholera toxin (100 ng/ml), and penicillin (50 U/ml) and streptomycin (50 μg/ml). Cell lines stably expressing biosensors were generated by retroviral or lentiviral infection, or by transfection with the PiggyBac transposase syste, (Yusa et al., 2011), followed by puromycin (1-2 μg/ml) selection and expansion of single clones. For each reporter, we isolated multiple stable clones with homogenous expression; all data reported in this study reflect representative behaviors that were highly consistent across all clones of each reporter (a minimum of three clones in each case).

For microscopy, we used a custom formulation with minimal background fluorescence, termed imaging-modified growth medium (iGM), which consists of DMEM/F12 lacking riboflavin, folic acid, and phenol red (Life Technologies). Lacking these three components had no effect on reporter kinetics, as indicated by experiments performed in normal growth medium lacking only phenol red (not shown), but allowed for more accurate quantification of reporter signals. iGM contained all supplements used in MCF10A culture medium (above), but with horse serum replaced by 0.1% (w/v) bovine serum albumin. In order to lower the amounts of extracellular pyruvate and facilitate measurements of cytosolic NADH-NAD^+^ redox (Hung et al., 2014) in some experiments, we used an alternate imaging medium formulation, termed iGM2, which consists of 95% DMEM (Life Technologies 31053) and 5% DMEM/F12 (Life Technologies 11039), supplemented with 0.3% bovine serum albumin, EGF, insulin, hydrocortisone, cholera toxin, and pen-strep as above. For Peredox calibration using lactate and pyruvate, DMEM (31053) with indicated lactate and pyruvate concentrations were made; cells were washed once to two times with the media prior to imaging. For GF titration, cells were placed in EGF/insulin-deficient medium for 2 days prior to imaging with appropriate concentrations of EGF and insulin. For glucose titration, DMEM (Life Technologies A14430) with indicated glucose concentrations were prepared (with residual F12 supplementation of 0.8% in media with 0.03 mM to 23 mM glucose); cells were washed two to three times with the respective media prior to imaging.

### Fluorescence microscopy

Time-lapse wide-field microscopy was performed as previously described (Hung et al., 2011; Albeck et al., 2013). Briefly, 1000-2500 cells were seeded 2-4 days prior in glass-bottom 24-well (MatTek) or 96-well plates (MGB096-1-2-LG-L; Matrical, Brook Life Sciences), with well bottom pretreated with a droplet of 5 μl type I collagen (BD Biosciences) to promote cell adherence. For experiments with drug addition, cells were placed in 240 μl imaging medium for 1-3 hours, until the addition of 60 μl as a 5x spike. Cells were maintained in 95% air and 5% CO2 at 37°C in an environmental chamber. Images were collected with a Nikon 20×/0.75 NA Plan Apo objective on a Nikon Eclipse Ti inverted microscope, equipped with a Lumencor SOLA or a Nikon Intensilight C-HGFI light source. Fluorescence filters were from Chroma: T-Sapphire (89000 ET Sedat Quad; or ET405/20x, T425LPXR, and ET525/50m), YFP (89002 ET ECFP/EYFP; or 41028), and RFP (49008 ET mCherry; or 41043 HcRed). With a Hamamatsu ORCA-ER or ORCA-AG cooled CCD camera, images were acquired every 5-8 min with 2×2 binning and an exposure time of 200-225 ms for T-Sapphire, 70-225 ms for YFP, 70-225 ms for RFP.

### Image Processing

Single cell traces were generated using the automated software DCellIQ (Li et al., 2010), followed by manual verification using a custom MATLAB program (MathWorks) to correct tracking errors, or using a custom MATLAB image processing pipeline (Sparta et al., 2015) using global optimization of cell tracks (Jaqaman et al., 2008). After background subtraction, DCellIQ or the MATLAB pipeline were used for image segmentation and tracking to determine nuclear masks based either on nuclear-localized Peredox-mCitrine or on the absence of YPet nuclear fluorescence of cytoplasmic-localized AMPKAR2. For a subset of the data, we additionally verified the automated tracking results manually. After cell tracking with the YFP images, the coordinates were applied to the other fluorescent channels. The nuclear masks were eroded by 1 μm to ensure the exclusion of cytoplasmic pixels; the nuclear T-Sapphire, CFP, YFP, and RFP signals were calculated as the mean pixel values within the nuclear masks in the respective images. The cytoplasmic CFP, YFP, and RFP signals were calculated as the mean pixel value within a cytoplasmic “donut” mask, which consisted of an outer rim 3-4 μm from the nuclear mask and the inner rim as the perimeter of the eroded nuclear mask or 2-3 μm from the original nuclear mask. NADH index was calculated as a ratio of the background-subtracted nuclear T-Sapphire to YFP signal. Akt index was calculated as a ratio of the background-subtracted nuclear RFP to cytoplasmic RFP signal. AMPK index was calculated as the ratio of the background subtracted cytoplasmic CFP to YFP ratio; because this ratio is linearly related to the fraction of unphosphorylated reporter molecules (Birtwistle et al., 2011), this signal was inverted by multiplying it by -1 and adding a positive value to set the lowest AMPK index in each experiment to approximately 0.

### Analysis and statistics of kinetics in reporter signals

A custom MATLAB algorithm was designed to identify peaks in the time-lapse signal of AMPK or NADH index within each cell. The AMPK or NADH index was first smoothed to remove spurious noise. Peaks and associated valleys in the index were identified by setting two local cutoff values, based on maximum and minimum values of the data within a sliding time window (typically 40 minutes). A peak was detected if both cutoff values were crossed by a rise and subsequent fall in the index. To define a “fluctuation score” for each cell, the amplitudes (difference between baseline and peak value) for all detected peaks were summed and normalized by the length of time the index was recorded. The fluctuation score for each cell thus increases with both the frequency and amplitude of peaks; examples of peak detection and corresponding scores are shown in Fig. S3A. Typically, >500 individual cell recordings were scored for each condition and plotted using a box plot, with the median shown as a red line, 25^th^-75^th^ percentile (the interquartile range) as the bottom and top of the box, respectively. Whiskers indicate the range of values falling within 1.5 times the interquartile range outside of the interquartile range; outliers beyond this range are not shown. Statistical comparisons between samples are displayed using notches on the box plots; two samples in which notches do not overlap differ in their medians at the 5% significance level.

### Replicates

Numbers of independent replicates are indicated in each figure legend as “n”; we define “independent replicate” here as a complete, separate performance of a time lapse imaging experiment with similar culture and treatment conditions, beginning from the plating of cells from bulk culture on an imaging plate and occurring on different days from other replicates. For all experiments shown, a minimum of 100 cells (not including daughters of cell divisions) were imaged and tracked for each condition. When possible, data from independent replicates were merged into a single data set using a set of calibration conditions for normalization of reporter signals; these datasets are indicated as “combined” in the figure legends. When calibration controls were not available, comparisons between conditions were made within replicate experiments, and we verified that similar trends were observable in every replicate; these datasets are indicated as “representative”. Unless noted otherwise, where single-cell recordings are shown, cells were chosen by random number generation in MATLAB with a threshold for minimum tracking time to eliminate cells in which recording was terminated prematurely due to failure of the tracking algorithm, and the chosen tracks were manually verified to be representative of successfully tracked cells and consistent with the overall range of cell behaviors. Cell recordings determined by manual inspection to have poor tracking or quantification accuracy were discarded.

## Acknowledgments

Imaging facilities were provided by the Nikon Imaging Center at Harvard Medical School and the Cell Biology Imaging Facility at UC Davis. We thank P. Tsou and L. Cantley for providing the AMPKAR plasmid; J. Coloff, G. Gao, J. Locasale, and T. Muranen for providing reagents; D. Clapham, T. Schwarz, V. Mootha, M. Vander Heiden, S. Gaudet, and members of the Albeck laboratory, Brugge laboratory, and the Yellen laboratory for their comments. This work was supported by a Stuart H. Q. and Victoria Quan predoctoral fellowship (to Y.P.H.), a U.S. Department of Defense Breast Cancer Research Program postdoctoral fellowship (W81XWH-08-1-0609 to J.G.A.), and the US National Institutes of Health (5-R01-CA105134-07 to J.S.B; R01 NS055031 to G.Y.).

---

**Figure S1.**
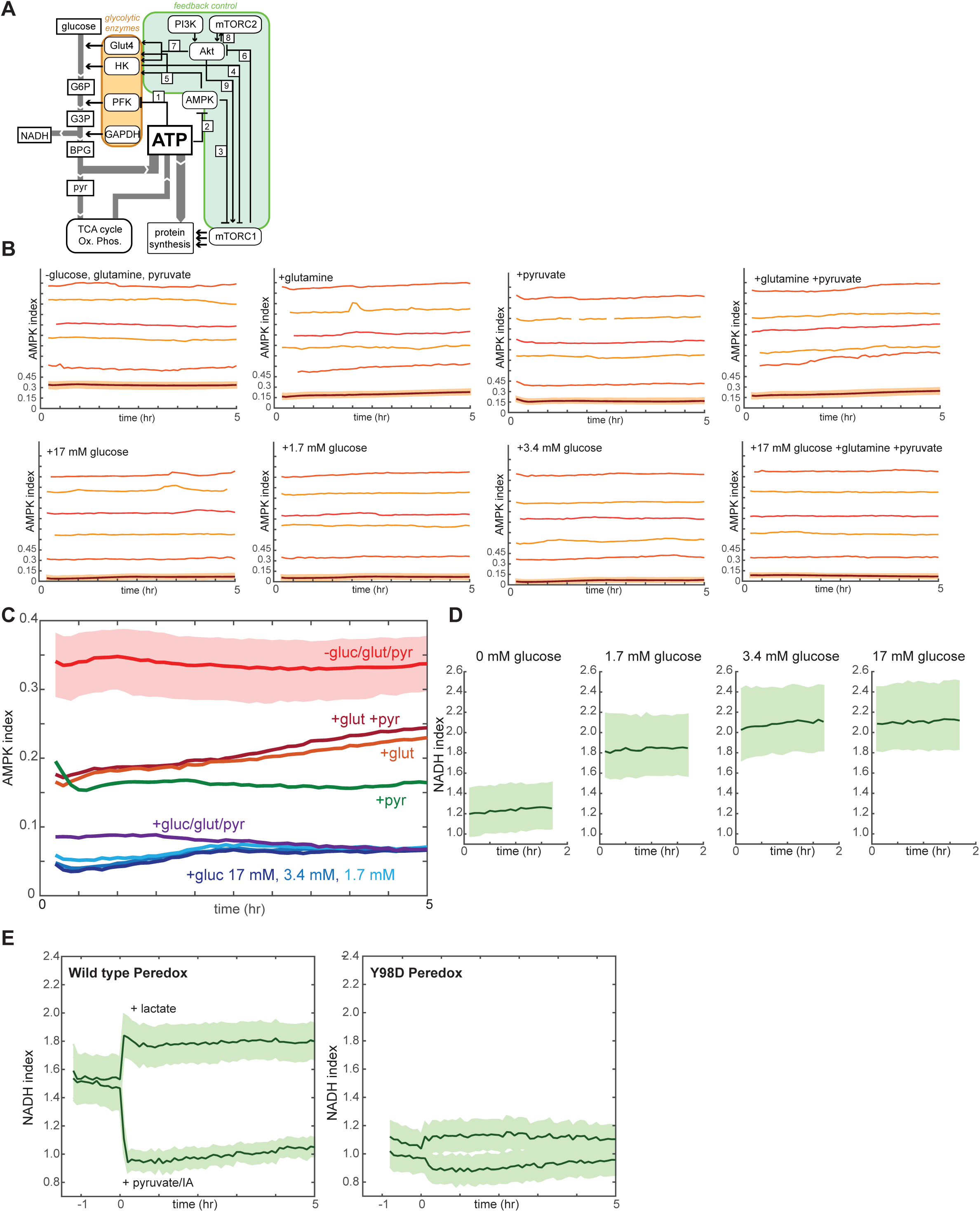
A. Schematic diagram of feedbacks connecting ATP production with glycolytic regulation. Flux of metabolites (white rectangles) is indicated by gray arrows, and glycolytic enzymes controlling this flux are grouped in the orange region. Known feedback connections are enclosed in the green region. These connections include: 1) suppression of PFK1 activity by ATP; 2) suppression of AMPK activity by ATP; 3) suppression of mTORC1 activity by AMPK; 4) suppression of mTORC1 by hexokinase (HK) during periods of low glucose flux; 5) stimulation of HK and Glut4 activity by AMPK; 6) inhibition of Akt by mTORC1; 7) stimulation of Glut4 and HK activity by Akt; 8) reciprocal positive regulation between Akt and mTORC2; and 9) activation of mTORC1 by Akt. The net effect of these feedback connections is to actively increase flux through glycolysis when ATP levels are low, as well as to actively suppress flux through glycolysis when ATP levels are high. B,C. AMPK index measurements in MCF10A cells cultured with various nutrients. Immediately prior to imaging, growth medium was replaced with iGM (see Methods) lacking pyruvate, glucose, and glutamine and supplemented with the indicated carbon sources. While long-term culture in the absence of all 3 nutrients led to cell death, measurements were possible for at least 5 hours. Mean and representative single cells are shown in (B). An overlay of the mean AMPK index for each condition, with the 25th-75th percentile range shown only for the minimal condition to allow for clarity (C). D. NADH index measurements in MCF10A cells cultured in the absence of pyruvate, glutamine, and serum, in the presence of the indicated glucose concentration. E. Lack of response of a mutant control NADH reporter to stimuli known to control cellular NADH levels. MCF10A cells expressing Peredox (left) or Peredox Y98D (right) were cultured in growth medium lacking pyruvate, and either lactate or a combination of pyruvate and IA were added at time 0. While the wild type reporter signal is rapidly increased by lactate and decreased by pyruvate/IA (left), the Y98D mutation in the NADH binding site of Peredox responds with reduced magnitude.

**Figure S2.**
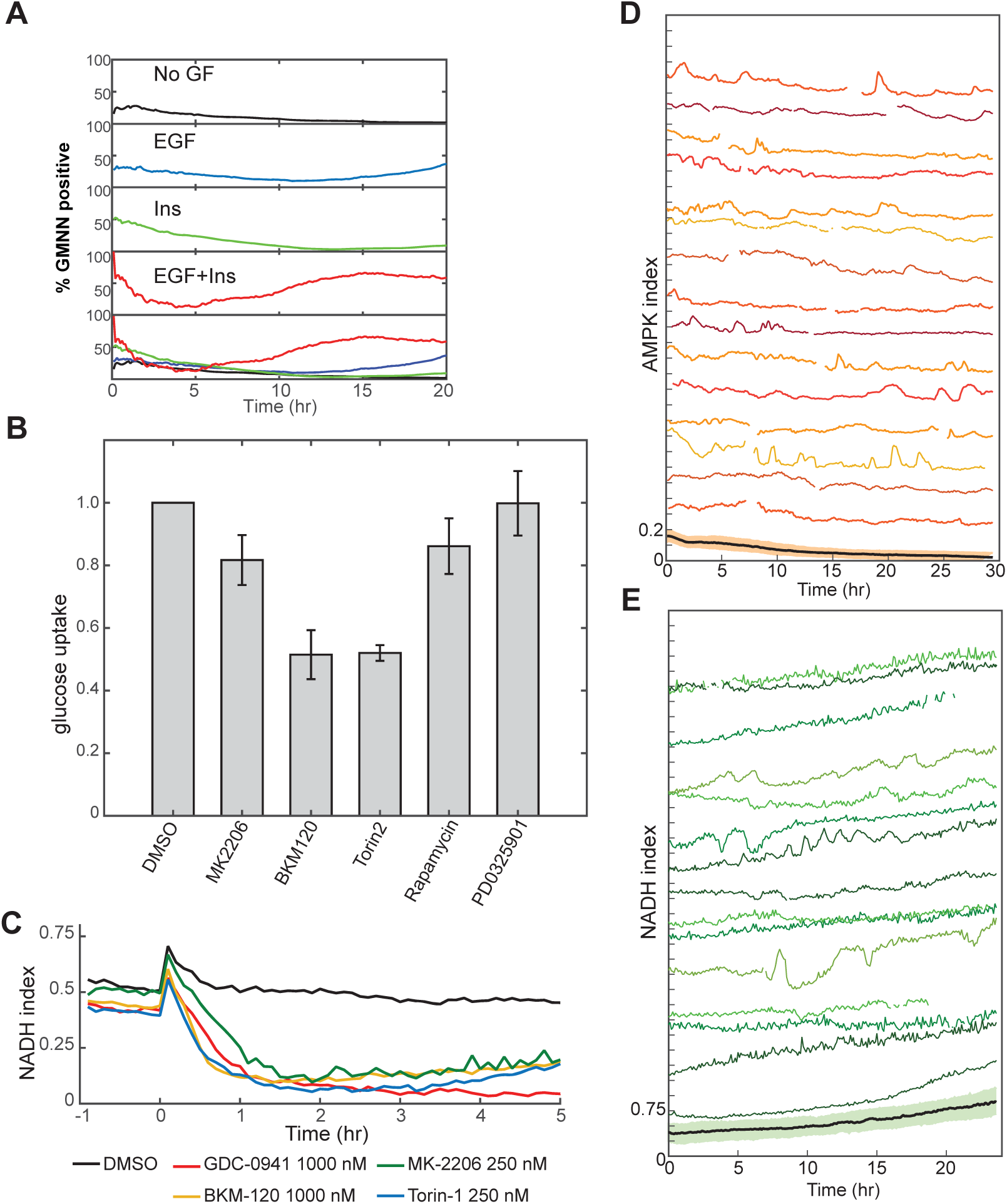
A. Proliferative index, calculated as the fraction of Geminin-mCherry-positive cells, monitored by live-cell microscopy in MCF10A cells under stimulation by the indicated GFs. Growth medium was iGM containing 3.4 mM glucose and 2.5 mM glutamine, and lacking pyruvate. B. Decrease in glucose uptake upon Pi3K/Akt inhibition. Glucose depletion from the medium was assayed immediately following a 2 hour period during which the cells were exposed to the indicated conditions. All measurements were made in the presence of EGF and Insulin, and measurements are normalized to the DMSO condition. C. Mean NADH index, measured by Peredox, following inhibition of the PI3K/Akt pathway. Prior to imaging, cells were placed in medium containing EGF and insulin. Inhibitors were added at time 0. D, E. Single cell variability in NADH and AMPK indices. MCF10A cells growing in iGM containing EGF and lacking insulin. Fifteen randomly selected individual cell traces and the population mean (bottom) are shown for each reporter, displaying sporadic peaks and fluctuations within individual cells.

**Figure S3.**
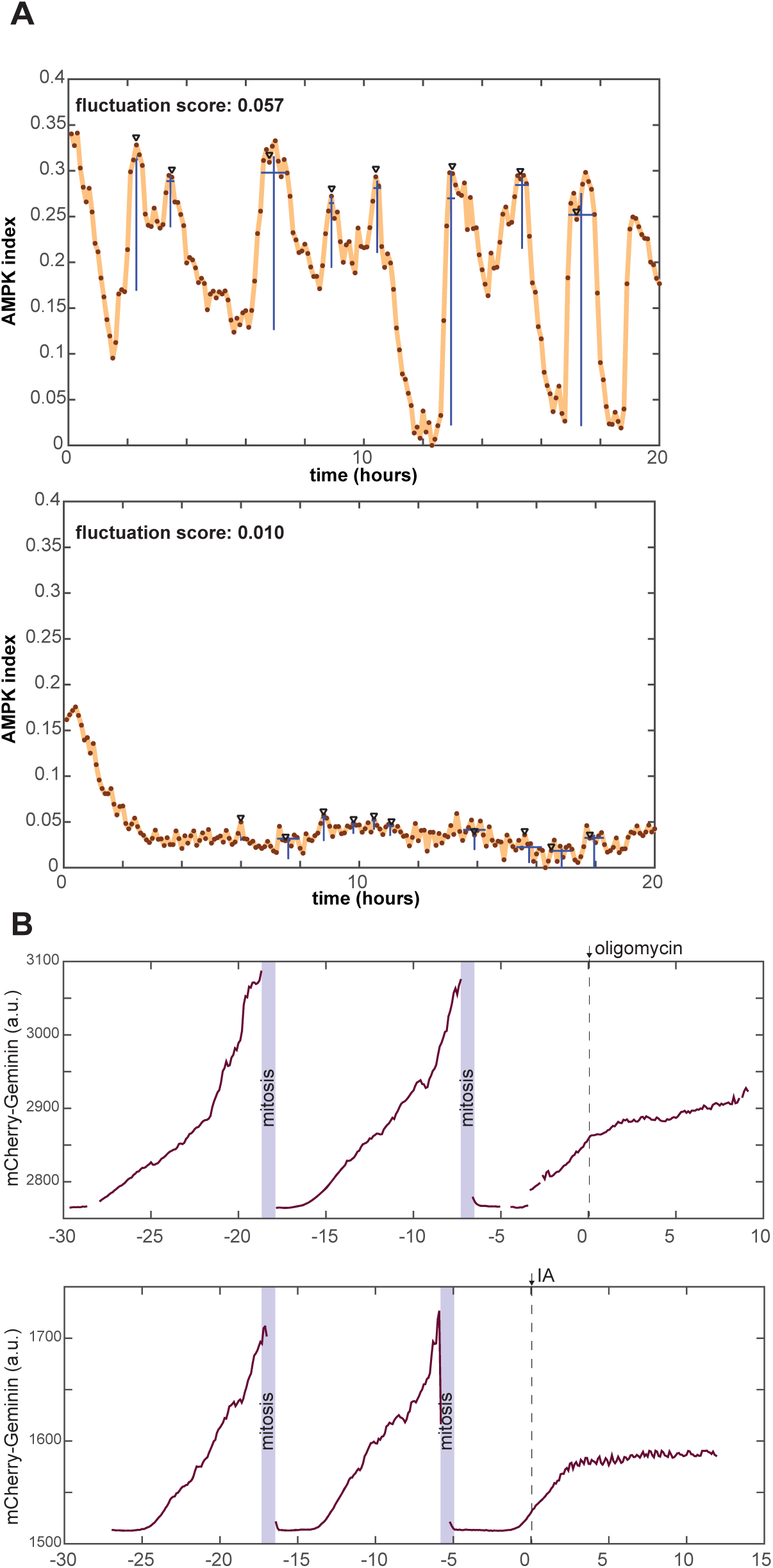
A. Identification of peaks and quantification of fluctuation trends in time-dependent reporter signals. AMPK index measurements from two cells are shown as examples (top: a combination of EGF and oligomycin, bottom: insulin). Peaks in each signal identified by an automated peak-finding algorithm (described in detail in the Methods section) are denoted by black triangles. Blue lines below each peak indicate the detected amplitude and duration for each peak. The fluctuation score for each cell is calculated by summing the amplitude for all peaks during the time period of measurement, and then dividing by the length of the time period in hours. B. Cell cycle responses during treatment with metabolic challenges. Representative traces for mCherry-Geminin fluorescence in individual cells first cultured in iGM and then treated either with oligomycin (A) or IA (B). In both cases, the imaged cell has progressed through 2 normal cell cycles prior to the time of treatment and has entered a third cell cycle prior to the time of treatment. In each case, cell cycle progression slows following drug treatment. In IA-treated cells, nearly all cells showing this form of arrest never complete mitosis before dying, while some oligomycin-treated cells do reach mitosis following the delay (Supplemental Movie 4).

**Figure S4.**
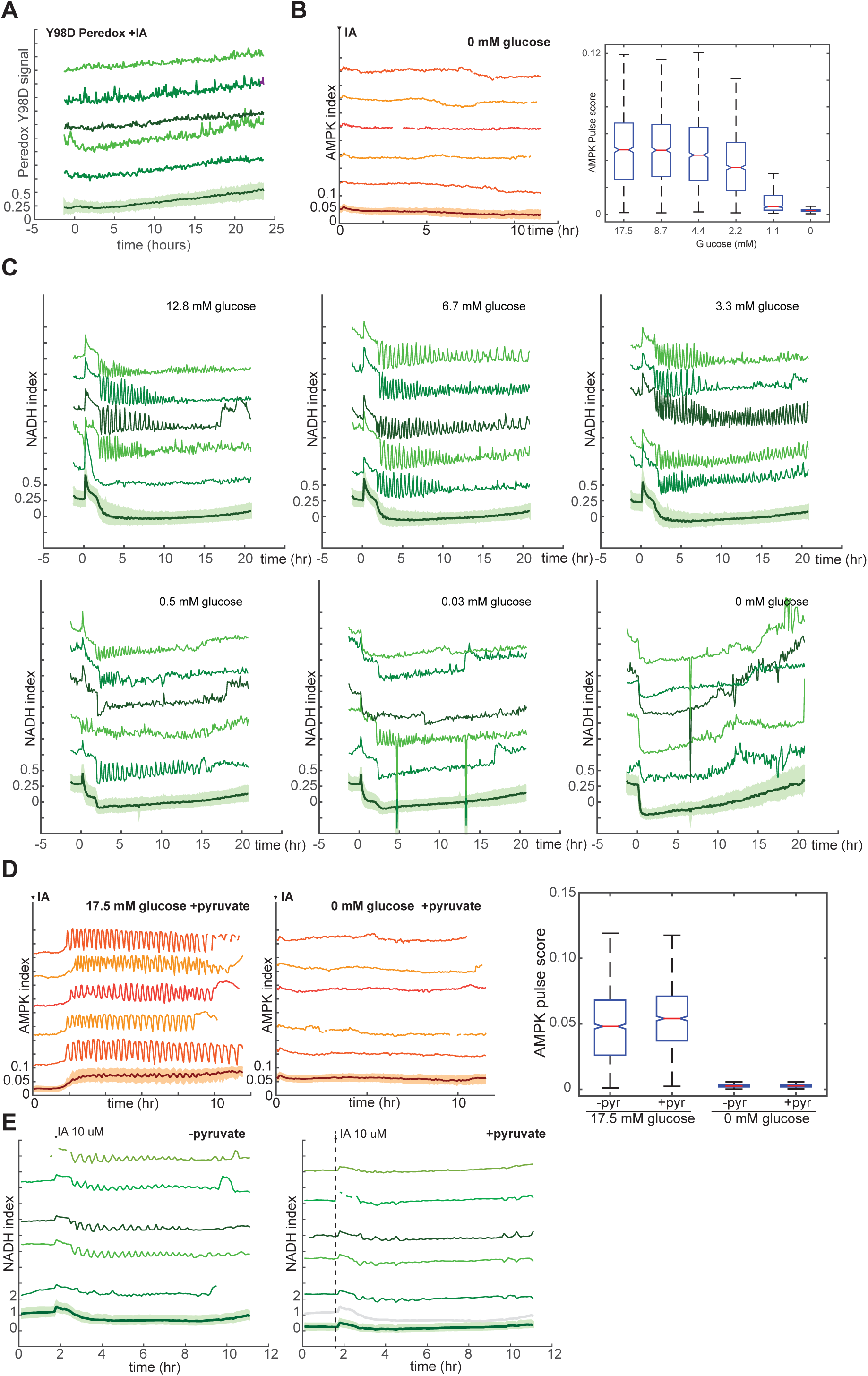
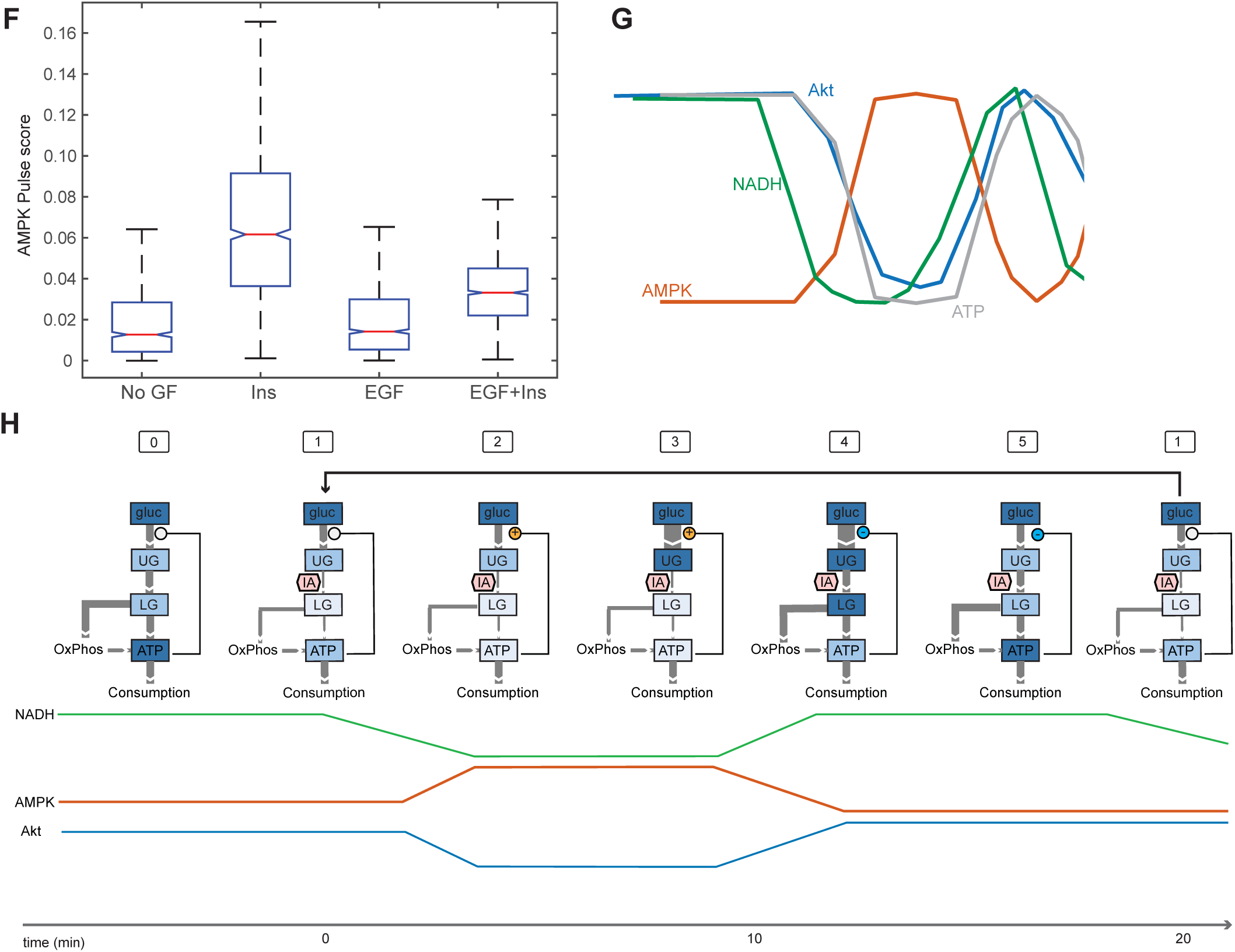
A. Lack of IA-induced oscillations in mutant Peredox Y98D control reporter. MCF10A cells expressing the binding site mutant NADH reporter were treated at time 0 with 10 μM IA in the presence of glucose (see Fig. 4D for the corresponding response of the wild type reporter). With the mutant reporter, regular oscillations cannot be detected, indicating that flucutations in NADH, and not other cellular changes affecting reporter fluorescence, are responsible for the observed oscillations in Peredox signal. B. Sensitivity of IA-induced AMPK index oscillations to glucose concentration. MCF10A cells expressing the AMPKAR2 reporter were cultured in the presence of glutamine and the indicated concentrations of glucose, treated with 10 μM IA at time 0. C. Sensitivity of IA-induced NADH index oscillations to glucose concentration. MCF10A cells expressing the Peredox reporter were cultured in the absence of pyruvate and presence of glutamine and the indicated concentrations of glucose, treated with 10 μM IA at time 0. D. Insensitivity of IA-induced AMPK index oscillations to pyruvate. MCF10A cells expressing AMPKAR2 were treated with 10 μM IA in the presence of glucose and pyruvate (left) or pyruvate alone (right).Pulse quantification is shown on the right, indicating that pyruvate does not alter oscillation behavior in the presence of glucose and does not provide fuel for oscillations in the absence of glucose. E. Absence of detectable NADH oscillations in the presence of pyruvate. MCF10A cells expressing Peredox were treated with 10 μM in the absence (left) or presence (right) of 50 mM pyruvate. The baseline NADH index is reduced in pyruvate-exposed cells (gray line in right plot-bottom shows mean NADH index in the absence of pyruvate for comparison), and IA-induced oscillations are not detectable (right). F. Pulse analysis of AMPK index in cells treated with 10 μM IA in the presence of the indicated GFs. G. Inferred relationship between ATP, NADH, AMPK activity, and AKT activity during oscillations. ATP is shown as the inverse of AMPK signal, indicating a phase shift relative to NADH (as measured by Peredox). This shifted relationship may contribute to the induction of oscillations, which often result from feedback in the presence of a delay. H. Model for oscillations in glycolysis and signaling upon inhibition of GAPDH by IA. Rectangles indicate pools of metabolites, with the cellular concentration of the pool indicated by the intensity of blue coloring (blue high, white low). Flux between pools is indicated by gray arrows, with rate of flux indicated by the width of the arrow. In state (0), cells in the presence of insulin and glucose operate at a high level of glycolytic flux and ATP levels are high. Upon IA treatment (1), flux from upper glycolysis (UG) to lower glycolysis (LG) is reduced, and ATP levels begin to fall. (2) Low ATP levels stimulate multiple positive feedback regulators of flux to UG (see Fig. S1A above). (3) Positive feedback drives an increase in the availability of glucose and UG metabolites; the decrease in Akt activity during this phase may be due to relief of mTOR inhibition by flux through HK (connection #4 in Fig. S1A), leading to negative feedback inhibition of Akt (connection #6 in Fig. S1A). (4) High levels of UG increase the levels of LG metabolites, overcoming the partial inhibition of flux imposed by IA; ATP levels rise, and net feedback switches from positive to negative. (5) Reduction of glycolytic flux leads to a relaxation of the levels of glycolytic metabolites; negative feedback deactivates. In a normal (non-IA-inhibited) state, this level of flux would be sufficient to maintain ATP levels, and the cell would return to a normal state (0). However, since IA remains present, flux from UG to LG is again inhibited and the cell returns to state (1), beginning a new repetition of the cycle.

**Figure S5.**
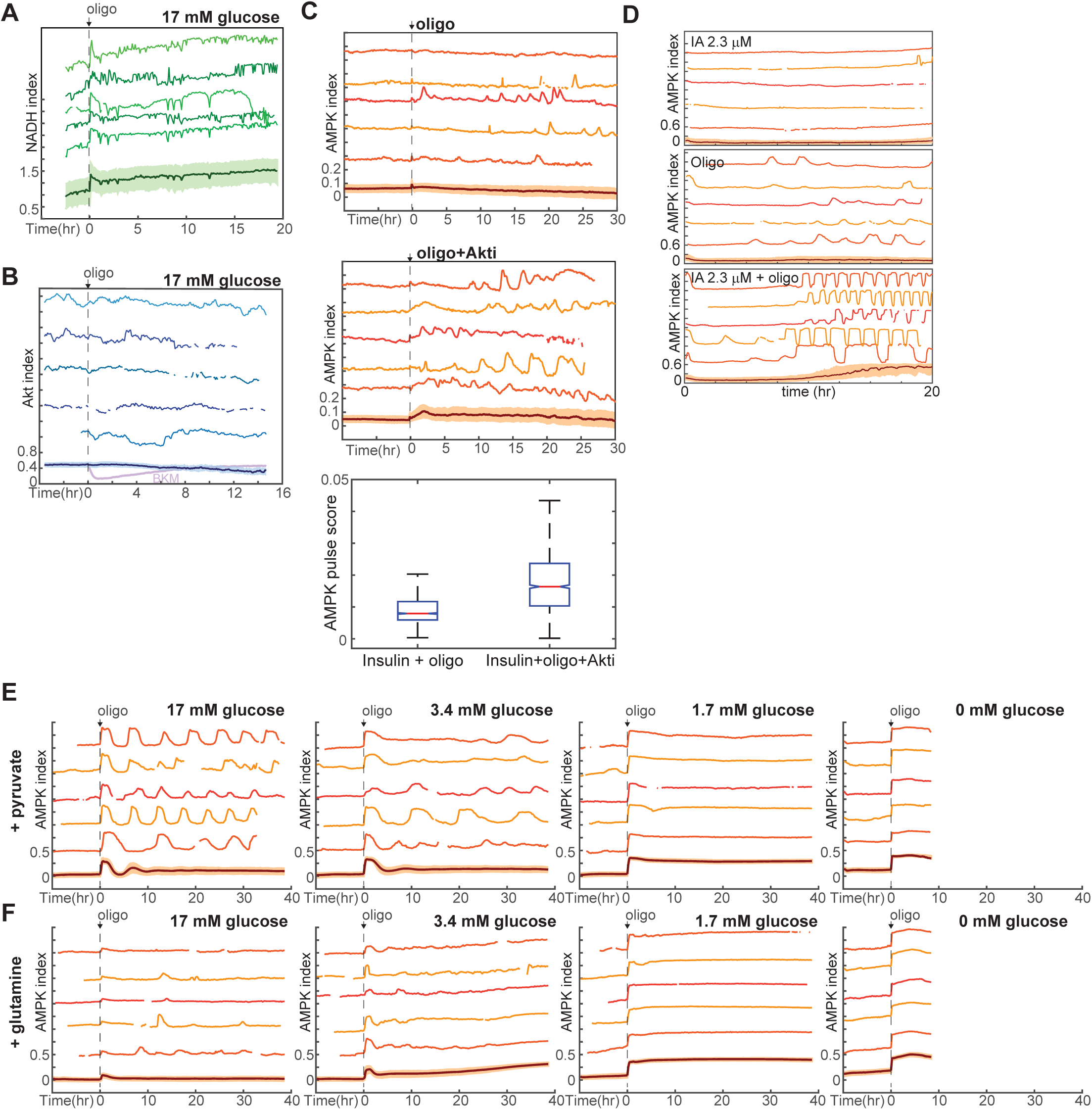
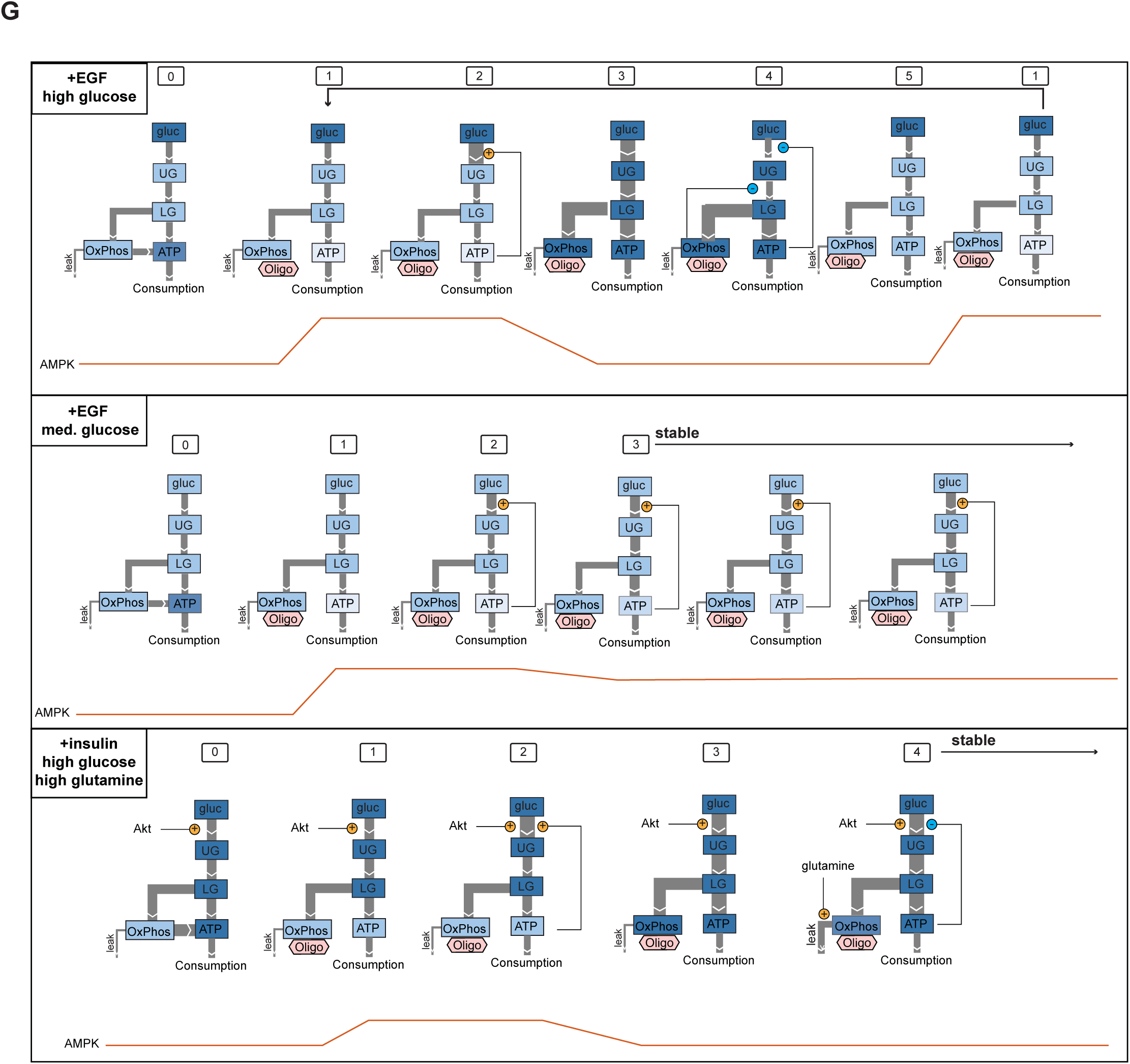
A. NADH index measurements in MCF10A cells expressing Peredox, in response to 1.8 μg/ml oligomycin. Cells were maintained in iGM lacking glutamine and pyruvate, containing 17 mM glucose. B. Akt index measurements in MCF10A cells expressing AKT-KTR, in response to 1.8 μg/ml oligomycin. Akt activity does not respond detectably to oligomycin treatment; inhibition in response to BKM-120 is shown in violet in the mean measurement (bottom) for comparison. C. Single-cell measurements of AMPK index in the absence or presence of an Akt inhibitor, treated with oligomycin at time 0. Cells were maintained in the presence of insulin and 17 mM glucose. D. Single-cell measurements of AMPK index in cells treated with a low concentration of IA, oligomcyin, or both. Medium included pyruvate, glutamine, and glucose; inhibitors were added at time 0. E,F. Dynamics of oligomycin-stimulated AMPK pulses in the presence of pyruvate or glutamine. Measurements of AMPK index were made as in Fig. 5A, but in the presence of 50 mM pyruvate (E) or 2.5 mM glutamine (F). Measurements under 0 mM glucose are truncated due to cell death that began approximately 5 hours following oligomycin treatment. While pyruvate does not alter the kinetics of pulsing, glutamine attenuates the pulse behavior. This effect of glutamine is not likely to be mediated by glutaminolysis, since the blockade of mitochondrial ATPase by oligomycin would prevent the majority of ATP production by this pathway. Another potential explanation is an acceleration of glycolysis by glutamine, as has been observed in some systems. G. Model for AMPK oscillations induced by oligomycin. We propose sequences of events that are consistent with three key scenarios found in our experimental data, one in which oligomycin triggers persistent oscillations (top), one in which cells reach stable adaptation at a lower level of ATP (middle), and one in which cells reach stable adaptation at a high level of ATP (bottom). Top panel shows the condition of EGF-stimulated cells in high glucose, which display strong oscillations upon oligomycin treatment (see Fig. 5A, first panel). In state (0), glucose metabolism operates at a moderately high level, and ATP levels are high. (1) When oligomycin is added, ATP production by oxidative phosphorylation is blocked, and ATP levels begin to fall; AMPK activity increases. (2) Positive feedback increases the rate of flux into upper glycolysis. (3) Increased glyocolytic flux increases ATP production and AMPK activity is reduced; pools of TCA intermediates in the mitochondria become saturated. (4) Negative feedback regulation of glycolysis is triggered both by high ATP levels and by buildup of citrate in the TCA cycle, leading to reduction of glycolytic flux. (5) ATP levels fall as a result of reduced glycolysis; TCA saturation dissipates through usage of TCA intermediates in other pathways and proton gradient leakage (together indicated as “leak” flux). In this state, glycolytic flux is insufficient to maintain ATP levels in the continuing presence of oligomcyin, and the cell returns to state (1). The middle panel indicates the condition of EGF stimulation in the presence of intermediate levels of glucose (∼2 mM; see Fig. 5A, third panel). (0) External glucose levels are not limiting in the basal state, and cells are able to maintain moderately high ATP levels. (1) Oligomycin blocks ATP production by oxidative phosphorylation, and ATP levels fall. (2) Positive feedback regulation increases the rate of flux through glycolysis; however, relative to the top panel, lower available glucose levels limit the extent to which glycolytic flux can increase. (3) Increased glycolytic flux leads to a moderate rise in ATP levels, but this increase is smaller than in the top model, and negative feedback to glycolysis is not triggered. This state persists stably as glycolysis continues to operate at a rate sufficient to supply some ATP in the continuing presence of oligomycin, but not enough to return to initial levels. The bottom panel shows the condition of insulin-treated cells in the presence of high glucose and glutamine (see Fig. 5D, lower right panel). In state (0), high Akt activity induced by insulin stimulates a high level of glycolytic flux and high ATP levels. (1) When oligomycin is added, ATP levels drop moderately, but not as far as in the top panels due to the higher glycolytic flux; AMPK activity increases moderately. (2) Positive feedback due to the loss of ATP stimulates further increase in glycolytic flux. (3) Increased glycolytic flux replenishes ATP and increases pools of TCA metabolites. (4) Negative feedback to upper glycolysis is triggered by an increase in ATP, but is counteracted by the insulin-stimulated high activity of Akt; glycolytic flux remains high. Meanwhile, glutamine entering the TCA cycle through anaplerotic reactions alleviates the negative feedback by allowing citrate to continue through the TCA cycle and preventing its buildup. Therefore, cells continue to be able to maintain a high level of glycolytic flux, providing high levels of ATP even in the absence of oxidative phosphorylation.

## Supplemental Movies

**Movie S1. Induction of AMPK oscillations by inhibition of glycolysis.** MCF10A-AMPKAR2 cells were imaged in iGM as in Fig. 3A, treated at time 0 with 10 μM IA. Color bar on upper right indicates AMPKAR signal, with white representing low activity and orange/brown representing high activity.

**Movie S2. Fluctuating AMPK activity in response to oligomycin.** MCF10A-AMPKAR2 cells were imaged in iGM as in Fig. 3B, treated at time 0 with 4.5 μg/ml oligomycin. Color bar on upper right indicates AMPKAR signal, with white representing low activity and orange/brown representing high activity.

**Movie S3. Continuous AMPK activity in response to CCCP.** MCF10A-AMPKAR2 cells were imaged in iGM as in Fig. 3C, treated at time 0 with 1.6 μM CCCP. Color bar on upper right indicates AMPKAR signal, with white representing low activity and orange/brown representing high activity.

**Movie S4. Cell fates in response to metabolic perturbation.** MCF10A-AMPKAR2-mCherryGMNN cells were imaged in iGM and treated with 10 μM IA, 4.5 μg/ml oligomycin, or 1.6 μM CCCP. The left column of images shows the YPet channel of AMPKAR as a cytosolic marker (yellow) and mCherryGMNN (red); the right column shows mCherryGMNN alone.

**Movie S5. Akt activity oscillations detected by AKT-KTR in response to IA.** MCF10A-AKT-KTR cells were imaged in iGM2 as in Fig. 4E, and were treated at time 0 with 10 μM IA.

**Movie S6. NADH activity oscillations detected by Peredox in response to IA.** MCF10A-Peredox cells were imaged in iGM2 as in Fig. 4D, and were treated at time 0 with 10 μM IA.

**Movie S7. AMPK activity dynamics as a function of glucose level.** MCF10A-AMPKAR2 cells were imaged in iGM modified to lack glutamine and pyruvate and to contain the indicated concentrations of glucose. Oligomycin (9 μg /ml) was added at time 0.

## Supplemental Files

MATLAB files used to analyze peak size and frequency in reporter signals are included in the compressed directory “AnalysisFiles”. For all analyses, the function ct_pulseanalysis was used with the parameters ‘narm’ set to 3 and ‘smooth’ set to 6. Subfunctions ct_getpeaks and ct_filter are required for ct_pulseanalysis, as is the MATLAB Signal Processing Toolbox.

